# Bifunctional Monosaccharides Preferentially Localize to Nuclear Subcompartments

**DOI:** 10.1101/2023.08.11.552905

**Authors:** Pavel Barahtjan, Juan M. Iglesias-Artola, Kristin Böhlig, Annett Lohmann, André Nadler

**Affiliations:** Max-Planck-Institute of Molecular Cell Biology and Genetics, Pfotenhauerstraße 108, 01307 Dresden (Germany)

## Abstract

Recent progress in glycan research has been driven by widespread implementations of metabolic oligosaccharide engineering. Complementing existing approaches, we here introduce bifunctional, UV-crosslinkable and clickable N-acetylglucosamine and N-acetylgalactosamine analogues, which enable direct visualization of the intracellular probe distribution as well as distinguishing monomeric and macromolecule-bound fractions. Using this feature, we find that monomeric N-acetylmonosaccharides partition into RNA-rich nuclear compartments such as nuclear speckles and nucleoli. This suggests the existence of spatially separated N-acetylmonosccharide pools within the nucleoplasm. Taken together, bifunctional N-acetylmonosaccharide probes are a powerful discovery tool for probing intracellular localization of monosaccharides and glycosylated macromolecules.

---

Glycan modifications of proteins and lipids are involved in crucial cell biological processes^1,2^, ranging from protein quality control in the endoplasmic reticulum^3^ to regulating ligand binding to cell surface receptors^4^. Genes encoding for glycosylation-related processes encompass 3-4 % of the human genome. In addition, 85% of the secretory proteins and the majority of nuclear and cytosolic proteins are estimated to be glycosylated^1^. Despite their obvious importance, detailed investigations of glycan function on the level of individual carbohydrate moieties have only become widespread after the implementation of metabolic oligosaccharide engineering (MOE) strategies^5^, an approach pioneered by the Bertozzi laboratory^6–8^. MOE approaches crucially rely on modifying monosaccharides with small functional groups, which are well-tolerated by the cellular glycosylation machinery^9–14^. The most prominent examples are azide- and alkyne-modified reagents^7,15^, which are usually administered to cells in acetate-masked form. After removal of the acetate groups by intracellular esterases, these probes are incorporated into protein and lipid-bound glycan structures and into carbohydrate polymers by cellular glycosyltransferases^5^. Incorporation of chemically modified MOE reagents can be increased by engineering enzymes in the early steps of the glycosylation pathways^16^. Key examples are the pyrophosphorylases AGX1/2 which generate nucleotide-sugar intermediates, substrates for many glycosyltransferases^17^. Ultimately, such approaches result in macromolecular glycan structures decorated with azide or alkyne groups. These functional handles enable specific derivatization by biorthogonal copper-mediated or strain-promoted azide-alkyne cycloaddition reactions for various analytical techniques^5^. Typical examples include facilitated mass spectrometric assignment of glycan structures in glycoproteomic approaches^18^, identification of glycan-interacting proteins^19^ and visualization of cell-surface glycans by fluorescence microscopy^20,21^. So far, most MOE approaches for visualizing glycans have focused on macromolecular structures while probes which also allow for analyzing intracellular localization and uptake rates of soluble monosaccharides on the single cell level by fluorescence microscopy are scarce. The primary reason for this methodological gap lies in the fact that introducing bulky fluorophores into small metabolites affects their function, thus rendering live-cell imaging approaches artefact-prone. In addition, soluble metabolic intermediates of minimally modified MOE reagents cannot be chemically fixed and are typically lost during cell permeabilization. We reasoned that monosaccharide probes bearing a minimal alkyne/diazirine linker^22^ would allow us to minimize these problems (Scheme 1).

**Scheme 1.**
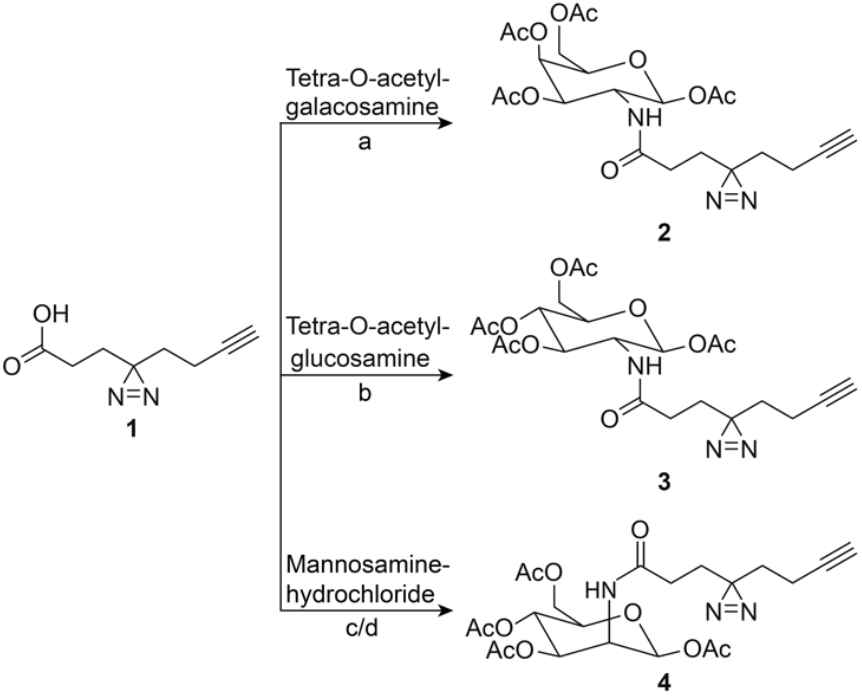
Synthesis of bifunctional, diazirine and alkyne containing N-acetylglucosamine, N-acetylgalactosamine and N-acetylmannosamine analogues **2**,**3** and **4**. a) HOBT, HBTU, DIEA, DMF, ON, RT, 48% b) HOBT, HBTU, DIEA, DMF, ON, RT, 52% c) HOBT, HBTU, DIEA, DMF, 3h, RT, 95% d) Ac2O, pyridine, ON, RT, 48%. The detailed synthesis procedure can be found in the supporting information (Chemical Synthesis, SI Scheme 1).

Since probes bearing either diazirine residues^9^ or combinations of diazirine groups and click handles^19,23,24^ have been successfully used for metabolic labelling of glycosylated proteins, we reasoned that this design should enable us to assess monosaccharide uptake rates and intracellular localization of soluble metabolites on the single cell level. Rapid photochemical crosslinking of the diazirine moiety should result in the formation of covalent protein-glycan adducts, which can be chemically fixed and stained by copper-mediated click chemistry for subsequent imaging via fluorescence microscopy, an approach that has been used in a limited number of examples for lipids^25–27^ but to the best of our knowledge not for mono- and oligosaccharides.

Here, we report the synthesis of bifunctional N-acetylglucosamine, N-acetylgalactosamine and N-acetylmannosamine derivatives. We used these probes to determine monosaccharide uptake and intracellular localization in different cell types by fluorescence microscopy. A comparison between +UV and -UV conditions enabled us to differentiate between small soluble species and chemically fixable macromolecules. Surprisingly, the N-acetylgalactosamine analogue (**2**) was found to accumulate in nuclear sub-compartments in all cell lines in its monomeric form, suggesting partitioning of small biosynthetic intermediates into metabolically highly active nuclear compartments.

The minimal, diazirine/alkyne modified linker **1** was synthesized according to a previously published protocol^22^. Tetra-O-acetylated glucosamine and galactosamine were then modified with **1** through HBTU/HOBT amide formation to yield probes **2** and **3** (Scheme 1). Probe **4** was synthesized by first coupling **1** with mannosamine-hydrochloride through HBTU/HOBT amide formation followed by acetylation (Scheme 1). We chose to apply the bifunctional probes in cell lines from different tissues and organisms to test the generality of the intracellular probe distribution. To this end, we used MDCK (a canine, epithelial cell line), HCT116 (a human, colorectal cancer cell line) and MIN6 (a murine, pancreatic beta cell line) cells^28^. The effect of the bifunctional monosaccharides on viability and metabolic activity was assessed via trypan blue cell staining and resazurin metabolic activity test. We found that the compounds had no effect on viability or metabolic activity (SI Figure 1). Cells were labelled with 50 μM of the respective probe for 10-360 min, followed by UV-cross-linking at 300 nm, cell fixation and fluorophore attachment via click chemistry (Figure 1a). The generated samples were imaged by confocal microscopy in multi-colour experiments utilizing ZO1 (for MDCK) and NaK-pump (MIN6 and HCT116) antibodies as plasma membrane markers and DAPI as a nuclear marker in addition to the monosaccharide stain. (Analysis of the probe characterization dataset is split between Figures 1 and 2 for greater clarity, exemplary images for each probe, time and cell line combination can be found in SI Figures 3-5).

**Figure 1.**
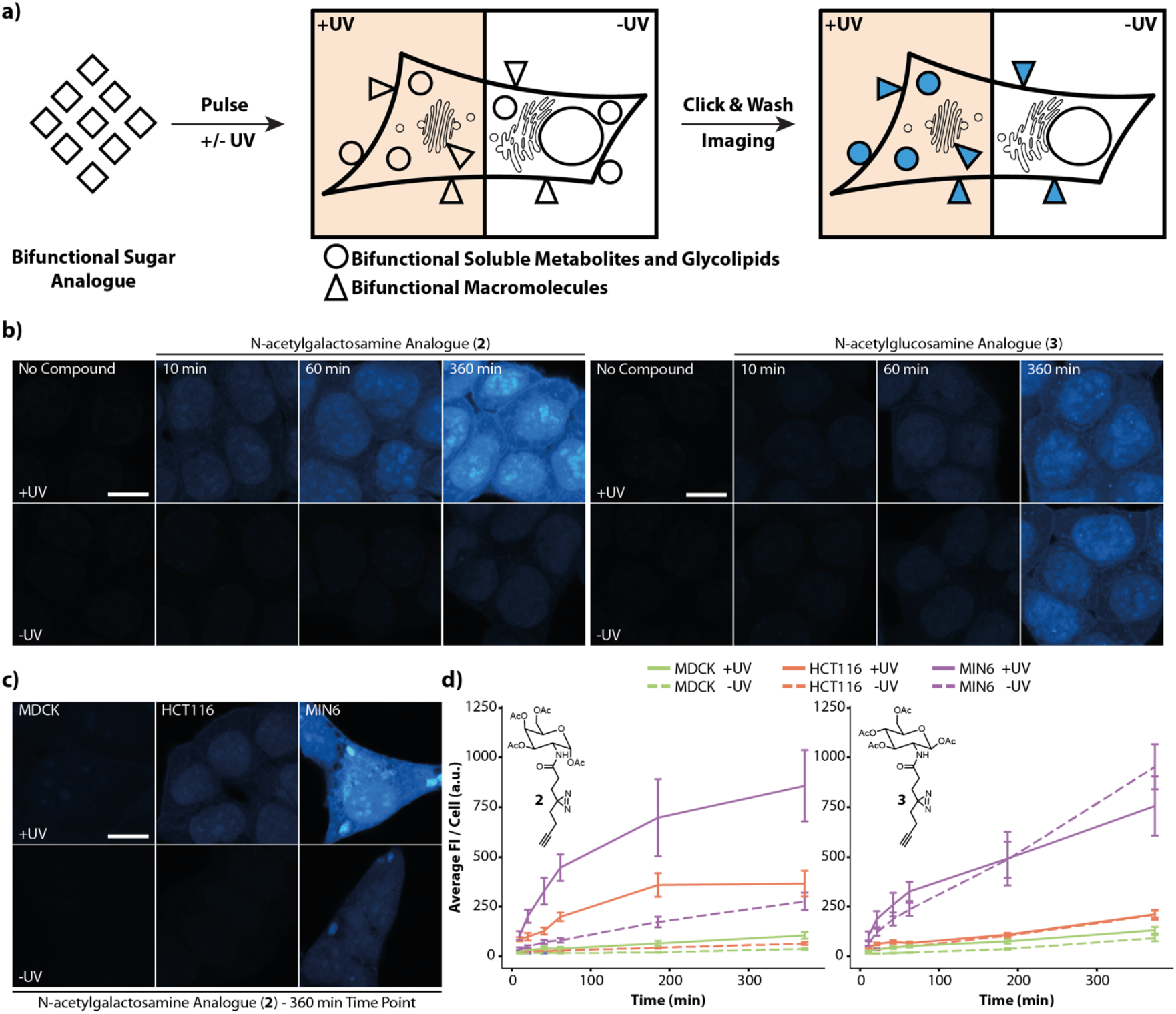
Bifunctional probe **2** persists as non-fixable metabolites while probe **3** is readily incorporated into cellular macromolecules. **a**) Experimental overview. Monosaccharides are fed to cells for different time periods. Afterwards, cells are either irradiated with 300nm UV light and fixed (+UV) or chemically fixed without irradiation (-UV). Subsequently, cells are clicked with the fluorophore and the intracellular localization is visualized by confocal microscopy. **b**) HCT116 cells were treated with 50 μM of probe **2** or **3** for defined time periods and afterwards processed as described in **a**). Selected time points are shown, see SI Figures 2-4 for complete time courses. **c**) Uptake of probe **2** in different cell lines after 360 min. The complete set can be found in SI Figure 2-4. **d**) Uptake dynamics of compounds **2** and **3** in different cell lines. Error bars indicate standard deviation of the mean. Fluorescence intensity changes over time were analysed using Python. Error bars indicate standard deviation. Scale bars = 14 μm. All images were acquired with the same microscopy settings. Images shown in **b**) were brightness-contrast adjusted in the same manner for better visualisation, the colour scale was defined using the 360 min probe **2** image. Solid and dashed lines are linear interpolations.

**Figure 2.**
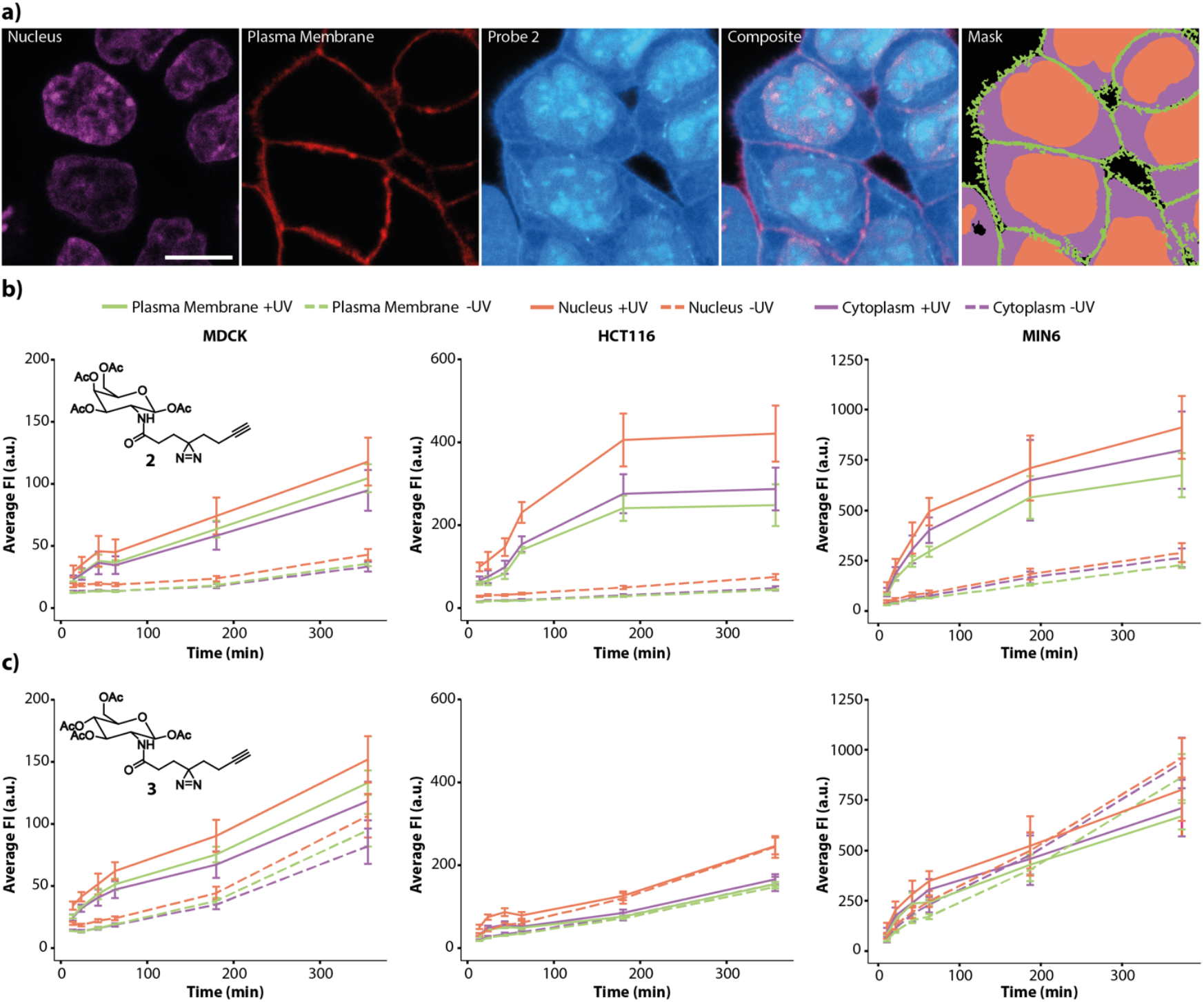
Subcellular distribution of compounds **2** and **3** in +UV and -UV conditions. Cells were segmented in three compartments: Plasma membrane, cytoplasm and nucleus **a**) Representative images of the markers (DAPI, NaK-Pump antibody) that were used to segment the cells, the probe channel and the resulting analysis mask. A +UV image from the 360 min time point of compound **2** in HCT116 cells is shown and brightness-contrast adjusted for visualization purposes. Scale bar = 14 μm. **b**,**c**) Quantification of subcellular localization of probe **2** (**b**) and probe **3** (**c**) by cell line over time. The analysed image dataset represents the same dataset that was used in Figure 1 and representative images for the complete time courses can be found in SI Figure 2-4. Error bars indicate standard deviation. Solid and dashed lines are linear interpolations.

We found that the utilized labelling protocol gave minimal background signal if untreated cells were used (Figure 1b, SI Figure 2), indicating that the observed fluorescence intensity pattern was indeed representative of the intracellular probe localization. In labelled cells, compound uptake increased over time for all cell lines (Figure 1b-d, SI Figures 3-5). For both probes, MIN6 cell labelling was significantly more pronounced compared to HCT116 and MDCK cells, suggesting very different uptake dynamics (Figure 1 c,d, SI Figures 3-5). In the case of the N-acetylgalactosamine analogue (**2**), irradiated (+UV) samples displayed much higher fluorescence intensities compared to non-irradiated samples (-UV), a trend that was consistently observed for all cell lines (Figure 1b,c, SI Figure 3-5). These data suggest that the predominant molecular species at all time points after labelling with **2** are small metabolites which cannot be chemically fixed with paraformaldehyde. In this case photochemical crosslinking is required to generate macromolecule-probe conjugates which in turn can be chemically fixed. Contrastingly, irradiated and non-irradiated samples generated by using N-acetylglucosamine analogue (**3**) show mostly negligible differences in fluorescence intensity suggesting that probe **3** is readily incorporated into fixable macromolecules (Figure 1b,d, SI Figures 3-5). We note, that the formation of covalent adducts can occur either via enzymatic glycosylation or via non-enzymatic S-glyco-modification^29,30^. While the observed plasma membrane stain is in line with enzymatic incorporation into glycosylated macromolecules (which was shown for structurally similar diazirine-containing N-acetylmonosaccharide probes)^31^, the later possibility cannot be ruled out.

We next assessed the subcellular probe distribution in a time dependent fashion. We observed incorporation of both probes into plasma membrane localized structures which is in line with exocytosis of glycosylated lipids and proteins derived from the secretory pathway (Figure 2, SI Figures 3-5). We also observed accumulation in the nucleus under all tested experimental conditions (Figure 2, SI Figures 3-5). In order to quantify the relative probe distribution patterns, we developed an automated image analysis pipeline using Python^32–34^ (see SI for details) that enables cell segmentation into plasma membrane, cytosolic and nuclear regions and a subsequent fluorescence intensity measurement (Figure 2a, see SI for details).

Using this approach, we found that the N-acetylgalactosamine probe **2** was incorporated with faster kinetics and to a larger overall amount in all analyzed compartments in HCT116 cells in +UV conditions compared with the N-acetylglucosamine probe **3** (Figure 2b,c). This contrasts with the situation in MIN6 cells, where faster uptake kinetics for **2** are maintained but overall incorporation levels after 6 hours are similar (Figure 2b,c). In MDCK cells, no major differences were observed for probes **2** and **3** in +UV conditions (Figure 2b,c). Most strikingly, we observed generally higher fluorescence intensities in nuclear regions compared to cytoplasmic and plasma membrane regions in all cell lines, indicating preferred localization of the N-acetylmonosaccharide probes in the nucleoplasm.

The similar fluorescence intensity -UV and +UV condition for the N-acetylglucosamine analogue **3** suggests a complete incorporation of the probe into proteins. In order to support the imaging approach and test for selectivity of the probes, we performed a biochemical assessment of probe incorporation into proteins by SDS-Page. We extended the set of probes by synthesizing the N-acetylmannosamine analogue **4** in order to cover a broader range of monosaccharide structures. Cells were treated with the respective compound for 6h and the fluorescent intensities were compared in the +UV and -UV conditions (Figure 3a). Compared to **2** and **3**, probe **4** showed a higher fluorescence intensity in the +UV condition but similar intensity under -UV conditions.

**Figure 3.**
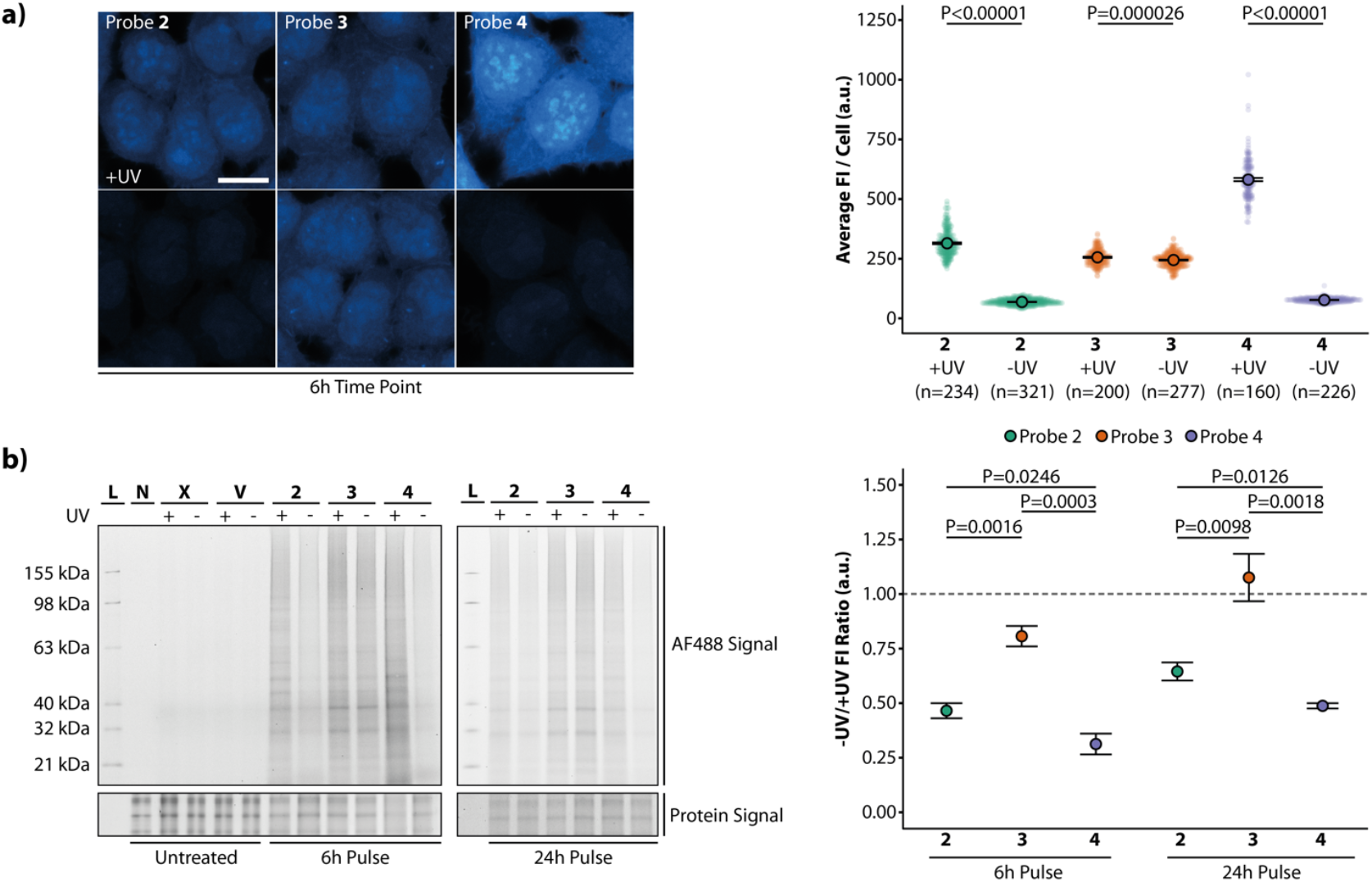
Selectivity and incorporation of monosaccharide analogues. **a)** Cells were treated with 50 μM of probe **2, 3** or **4** for 6h and afterwards processed as described in Figure 1. The average fluorescence intensity per cell was calculated with the same pipeline described before and is shown in the graph. Scale bar = 14 μm. Statistical analysis was performed using a one-sided Student’s t-test test. n represent the number of analysed cells per condition. Error bars indicate standard error of the mean. Images were brightness-contrast adjusted in the same manner for better visualisation, **b)** Cells were treated with 50 μM of probe **2, 3** or **4** for 6h and 24h and afterwards irradiated with 300nm UV light and lysed (+UV) or lysed without irradiation (-UV). The whole cell lysate was subjected to click labelling and proteins were analysed by SDS-Page (two-colour gel image). L = Ladder, N = No-click control, X = Untreated and V = Vehicle control. The right panel shows the fluorescence ratio of -UV and +UV. Statistical analysis was performed using a one-sided Students t-test test. Experiments were performed 4 times (all gels can be found uncut in SI Figures 7-8). Error bars indicate standard error of the mean.

This suggests lower incorporation of probe **4** into fixable macromolecules under the chosen experimental conditions. This finding is supported by SDS-Page results (Figure 3b). Cells were treated for 6h or 24h with the respective compound and afterwards irradiated with UV light and lysed (+UV) or lysed without irradiation (-UV). We used the ratio of -UV/+UV as a measure of incorporation efficiency which allows to calculate the percentage of covalently bound monosaccharide molecules. The N-acetylglucosamine analogue **3** was nearly completely incorporated into proteins (∼81% at 6h and ∼100% at 24h). The N-acetylgalactosamine analogue **3** showed a lower incorporation (∼49% at 6h and ∼65% at 24h) into proteins and the N-acetylmannosamine analogue **4** exhibited the lowest ratio (∼31% at 6h and ∼50% at 24h). These results are in line with the results from fluorescent microscopy.

Since the unexpectedly strong nuclear stain appeared to be non-homogeneous, we carried out a series of colocalization experiments with nuclear markers to identify intranuclear sites of N-acetylsaccharide analogue enrichment. To this end we imaged HCT116 cell samples that were treated with compounds **2** and **3** for 6 hours and subsequently counterstained with DAPI and markers for either nuclear speckles (NS), the dense fibrillar component (DFC) of the nucleolus, or Histone H3K9me3 (a heterochromatin marker). We found that both the DFC and the NS exhibited a stronger N-acetylsaccharide stain in both +UV and -UV conditions (Figure 4, SI Figure 6), compared to the remaining nucleoplasm, whereas the heterochromatin region did not show enrichment in any condition (Figure 4, SI Figure 6, See SI for image analysis details). These effects were even clearer in high-resolution experiments carried out by using the SoRa settings of an Olympus spinning disk microscope (Figure 4F). Specifically, we found 1.39 ± 0.01 (mean ± SE, also applies to following values) fold enrichment in DFC and 1.22 ± 0.01-fold enrichment in the NS for compound **2**. The respective values for **3** are 1.35 ± 0.01-fold for DFC and 1.23 ± 0.01-fold for the NS (Figure 4d,e). For the N-acetylgalactosamine analogue **2**, the increased signal in nuclear sub-compartments in +UV conditions suggests partitioning of small, not chemically fixable, metabolites since such molecules constitute the main contribution of the observed total fluorescence intensity (compare Figures 1,2).

**Figure 4.**
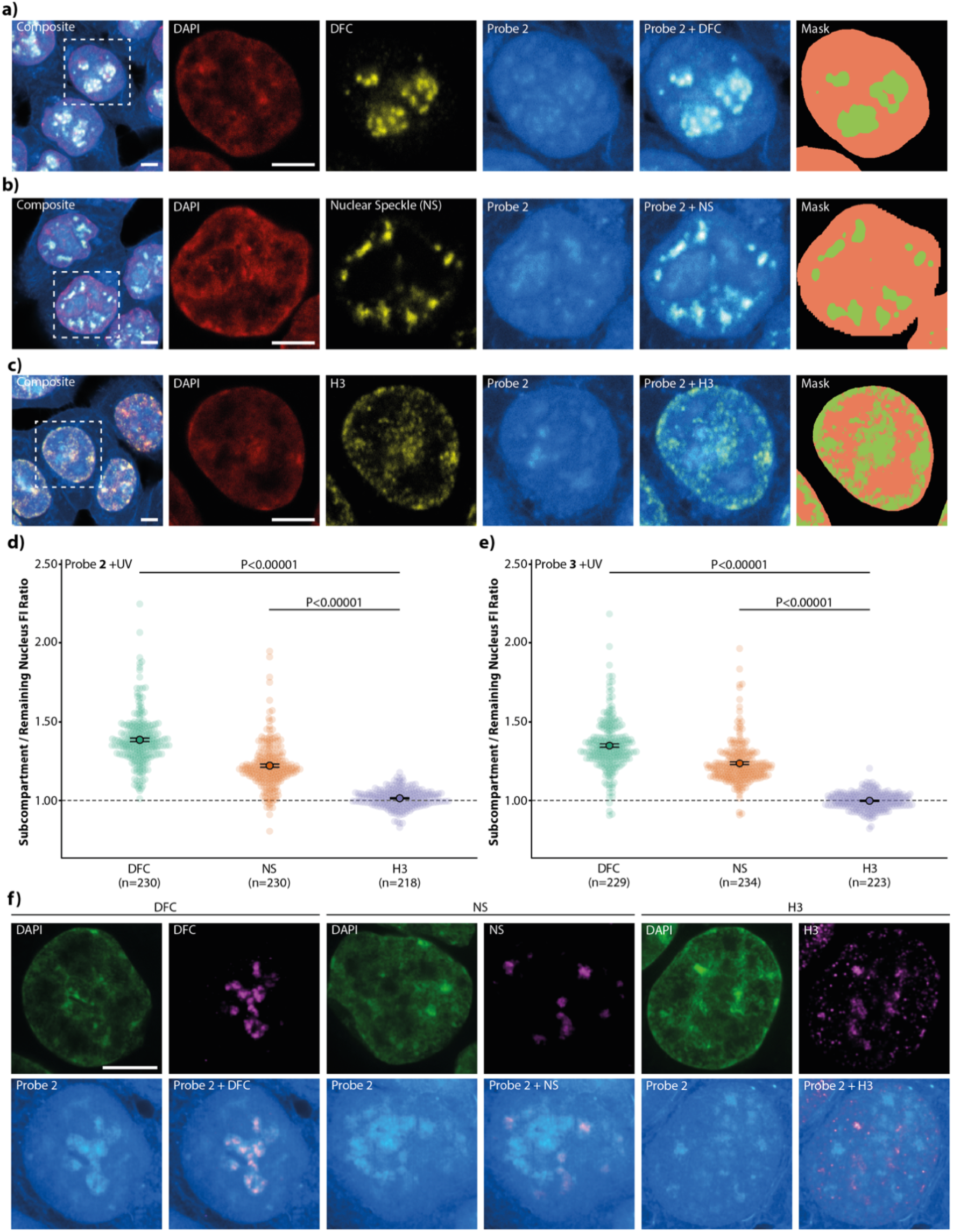
N-acetylsaccharides probes are enriched in nuclear subcompartments. **a**,**b**,**c**) Representative images displaying DAPI, the respective compartment marker (antibody stainings for DFC, NS and H3K9me3, abbreviated as H3 in the figure), their respective composites and the segmentation mask for quantification. Green areas in the masks indicate the compartment marked by the respective antibody, while the remaining nuclear area is marked in orange. Dashed squares in composite panels indicates a nucleus that is magnified for better visualization. Scale bars = 10 μm. **d**,**e**) Enrichment of probe **2** (**d**) and probe **3** (**e**) in nuclear subcompartments. Error bars indicate standard error. Statistical analysis was performed using a two-sided Mann-Whitney-U test. n Numbers represent the number of analysed nuclei per condition. Complementary images, composites and -UV plots can be found in SI Figure 6. **f**) High-resolution images displaying DAPI, the respective compartment marker and their respective composite. Scale bar = 5 μm. Images shown in **b**) were brightness-contrast adjusted in the same manner as above for better visualisation.

Intriguingly, all compounds appear to partition into metabolically highly active RNA-rich sub-nuclear structures, specifically the nucleolar DFC and the NS, which are the sites of RNA and protein processing. In the case of **2** and **4**, this clearly happens in the form of small, soluble species. Thus, it is possible to speculate that both the DFC and nuclear speckles are metabolic hotspots where glycosylation reactions are facilitated by higher local substrate concentrations. This notion is additionally supported by the finding that the nuclear N-acetylglucosamine kinase NAGK indeed localizes to nuclear speckles^35^. As of now, it is unclear which precise cellular mechanisms counteract rapid diffusion-driven equilibration of probe distribution throughout the nucleo- and cytoplasm. One intriguing possible explanation could be the presence of low-affinity, low-specificity sugar binding sites in scaffolding proteins of the respective nuclear compartments, which would result in a readily available supply of glycosylation substrates. Finally, our observation of high nuclear monosaccharide concentration specifically in nuclear compartments where covalent RNA modifications occur is well in line with the recent discovery of glycan-decorated RNAs^36^. Taken together, our bifunctional monosaccharide analogues function as discovery tools for imaging applications, especially with regard to determining the subcellular localization of small soluble biomolecules that are not retained by traditional chemical fixation methods.

## Acknowledgements

We kindly thank the Genome Engineering Facility of MPI-CBG for providing HCT116 cells, Dr. Alf Honigmann for providing MDCK cells, Prof Dr. Michele Solimena and Prof. Dr. Jun-ichi Miyazaki for providing MIN6 cells. Furthermore, we would like to thank the Light Microscopy Facility of MPI-CBG for their expert support. A.N. gratefully acknowledges financial support by the European Research Council (ERC) under the European Union’s Horizon 2020 research and innovation program (grant agreement no GA 758334 ASYMMEM). A. N. and P. B. gratefully acknowledge financial support by the Deutsche ForschungsgemeinschaY (DFG) via the TRR83 consortium.

## Supplementary information

Methods and supplementary figures are provided in an extra document.

Original imaging data are deposited in the following repository: https://dx.doi.org/10.17617/3.8i

Python scripts are accessible under: https://git.mpi-cbg.de/publications/nadler/2022-barahtjaniglesiasartola-monosaccharide-imaging

