## Supporting Information for "Bifunctional Monosaccharides Preferentially Localize to Nuclear Subcompartments"

#### Supporting Information - Material and Methods

##### Cell Culture

*HCT116* cells were cultured in McCoy's 5A GlutaMAX™ medium (36600-021, Life Technologies) supplemented with 10 % not-heat inactivated fetal bovine serum (10270-106, Life Technologies) and 100 $\mu$g ml$^{-1}$ antibiotic Penicillin-Streptomycin (10 000 U/mL, 15140-122, Life Technologies). Cells were trypsinised with 0.05 % trypsin (25300054, Life Technologies) and passaged after every 48h to 72h in a 1:10 ratio into Nunc EasYFlask 75 cm$^2$ Nucleon Delta Surface (156499, Thermo Scientific) flasks. Cells were grown at standard cell culture conditions (37 °C, 5 % $CO_2$).

*MDCK* cells were cultured in MEM GlutaMAX™ medium (41090036, Life Technologies) supplemented with 10 % heat inactivated fetal bovine serum (10270-106, Life Technologies), 100 $\mu$M non-essential amino acids (11140050, Life Technologies), 1 mM sodium pyruvate (11360070, Life Technologies) and 100 $\mu$g ml$^{-1}$ antibiotic Penicillin-Streptomycin (10 000 U/mL, 15140-122, Life Technologies). Cells were trypsinised with 0.25% trypsin (25200056, Life Technologies) and passaged after every 48h in a 1:10 ratio into Nunc EasYFlask 75 cm$^2$ Nucleon Delta Surface flasks (156499, Thermo Scientific). Cells were grown at standard cell culture conditions (37 °C, 5 % $CO_2$).

*MIN6* cells were cultured in DMEM High Glucose medium (11965092, Life Technologies) supplemented with 15 % heat inactivated fetal bovine serum (10270-106, Life Technologies), 70 $\mu$M 2-Mercaptoethanol (M6250, Merck) and 100 $\mu$g ml$^{-1}$ antibiotic Penicillin-Streptomycin (10 000 U/mL, 15140-122, Life Technologies). Cells were trypsinised with 0.25 % trypsin (25200056, Life Technologies) and passaged after every 72 h in a 1:10 ratio into Nunc EasYFlask 75 cm$^2$ Nucleon Delta Surface flasks (156499, Thermo Scientific). Cells were grown at standard cell culture conditions (37 °C, 5 % $CO_2$).

##### Preparation of Samples Labelled with Bifunctional Probes

Monosaccharide labelling experiments were performed using 96-well plates (655891, Greiner Bio-One). Cells were seeded one day prior to staining with 35000 cells/well. Stock solutions (10 mM) of monosaccharide analogues were prepared by dissolving in ethanol. The final labelling solutions were prepared by diluting the stock solution in the respective cell culture medium to reach a final monosaccharide analogue concentration of 50 $\mu$M. For the labelling experiments, the cell growth medium was removed and replaced by the sugar analogue containing medium. Cells were then incubated for the indicated time at 37 °C and 5% $CO_2$. At the defined time point, cells were washed twice with cell culture medium and either irradiated for 10 s using a 300 nm LED and then fixed (Laser Component GmBH) or immediately fixed without irradiation. Cell fixation was carried out removal of the medium and addition of 4 % paraformaldehyde (30525-89-4, TCI)(in PBS). The fixation solution was left on the cells for 20 min at room temperature. After fixation, cells were washed twice with paraformaldehyde-quenching and permeabilising solution (100 mM glycine, 0.3% Triton X-100 in PBS)(1.04201.1000, Merck / 37240.0, Serva) and afterwards placed in the quenching and permeabilising solution for 30 min at RT. Afterwards, cells were washed twice with PBS prior to click chemistry. Copper-mediated click chemistry was used to stain (macro)molecules containing an alkyne group. A click staining solution was prepared with 2 $\mu$M Azide dye (CLK-1276, Jena Bioscience), 5 mM sodium L-ascorbic acid (95210, Merck), 500 nM THPTA (CLK-1010, Jena Bioscience), 100 $\mu$M $CuSO_4$ (7758-98-7, ACROS Organics™) and 100 mM HEPES (pH 7.25). Before the application of the click solution cells were washed three times with 100 mM HEPES (pH 7.25). Subsequently, 100 $\mu$L of click staining solution were added per well for 20 min at 37 °C. Cells were clicked three times (without washing steps in between). To remove remaining free dye, the cells were washed three times with 0.1 % Triton X-100 (in PBS), the final wash was left for 30 min. Finally, the cells were subsequently washed three times with PBS and stored in PBS at 4 ºC prior to imaging.

##### Immunofluorescence Stainings of Monosaccharide Treated Cells

For the colocalization experiments, the click-chemistry labelled cells were further co-stained using different antibodies and DAPI as a nuclear marker. Before antibody application, blocking solution (0.1% Triton, 2% BSA (A8806, Sigma-Aldrich) in PBS) was added for 1h at RT. For the plasma membrane staining, two different antibodies were used. MIN6 and HCT116 cells were treated with a Sodium Potassium ATPase antibody (ab198367, abcam, diluted 1:100 in blocking solution) for 1h at RT. For MDCK cells, the plasma membrane was stained using a ZO-1 antibody (33-9100, Invitrogen, diluted 1:200 in blocking solution). For staining nuclear compartments, a fibrillarin antibody (ab5821, abcam) was used for the DFC of nucleoli, an antibody against SC-35 (ab11826, abcam) for nuclear speckles (both in 1:300 dilution) and an antibody against Histone H3K9me3 (39062, activemotif) as a heterochromatin marker in a 1:1000 dilution. All dilutions were performed in blocking solution. Primary antibodies were applied over night at 4°C. After staining, the cells were washed three times with PBS and stained with AlexaFluor647 (A32728 or A32733, ThermoFisher Scientific) (diluted 1:500 in blocking solution) for 1h at RT and subsequently washed 3 times with PBS. The nucleus was stained with 300 nM DAPI (D9542, Sigma-Aldrich) solution in PBS for 5 min at RT. Afterwards, cells were washed three times with PBS.

| **Primary/Secondary Antibodies** | **Type** | **Supplier** | **Catalogue#** | **Host species** | **Dilution** |
|---|---|---|---|---|---|
| Alexa Fluor® 647 Anti-Sodium Potassium ATPase | Monoclonal | abcam | ab198367 | Recombinant/Rabbit | 1:100 |
| ZO-1 | Monoclonal | Invitrogen | 33-9100 | Mouse | 1:200 |
| Fibrillarin | Polyclonal | abcam | ab5821 | Rabbit | 1:300 |
| SC-35 | Monoclonal | abcam | ab11826 | Mouse | 1:300 |
| Histone H3K9me3 | Polyclonal | activemotif | 39062 | Rabbit | 1:1000 |
| AlexaFluor647 | Polyclonal | ThermoFisher | A32728 | Goat (anti-mouse) | 1:500 |
| AlexaFluor647 | Polyclonal | ThermoFisher | A32733 | Goat (anti-rabbit) | 1:500 |

**Assessment of Cell Viability/Metabolic Activity**

*Trypan Blue Approach*: HCT116 cells were seeded one day prior to experiment into 24-well plates (142475, Thermo Scientific) with 100000 cells/well. On the next day, medium was removed and 1ml of medium containing the compound of interest was added to the cells and incubated for 6h or 24h. The medium was collected and 200 $\mu$L Trypsin (25300054, Life Technologies) was added to the cells and incubated for 3 min at 37 °C. Subsequently, 800 $\mu$L of medium was added and the cells were resuspended by pipetting up and down several times and the solution was added to the removed medium. 10 $\mu$L of cell solution was mixed with 10 $\mu$L of Trypan Blue (15250061, Life Technologies) and afterwards applied to cell counting chamber slides (C10228, invitrogen) and viability was determined using the Countess Automated Cell Counter (C10227, invitrogen).

*Resazurin Approach*: HCT116 cells were seeded one day prior to experiment into 96-well plates (89626, ibidi) with 30000 cells/well. On the next day, medium was removed and 100 $\mu$L of medium containing the compound of interest was added to the cells and incubated for 6h or 24h. After the respective time, 20 $\mu$L of resazurin assay buffer solution (ab112119, abcam) was added to the wells and incubated for 1h at 37 °C. Fluorescence intensity was measured with a BMG plate reader (Ex/Em = 540/590).

**Assessment of Incorporation of Monosaccharides into Proteins by SDS-Page**

Cells were seeded one day prior to protein extraction with 20000 cells/well into 96-well plates (655891, Greiner Bio-One). On the next day, medium was removed and 100 $\mu$L of medium containing the compound of interest (medium, vehicle without monosaccharides and the respective monosaccharides) was added to the cells and incubated for 6h or 24h. Monosaccharides were added to a final concentration of 50 $\mu$M. After the chosen time, cells were washed three times with ice-cold PBS and subsequently cells were UV irradiated for 10s and 200 $\mu$L of ice-cold RIPA buffer (89900, Thermo Scientific) was added immediately to the cells. For non-irradiated cells, RIPA buffer was directly added after washing. Protein solution was centrifuged (20000 x g, 15 min) and concentrated using Amicon Ultra 3 kDa centrifugal filters (UFC800324, Merck Millipore). Protein concentration of cell lysate was determined using BCA Protein Assay Kit (23225, Thermo Scientific).

Cell lysate was clicked with 2 $\mu$M Azide dye (CLK-1276, Jena Bioscience), 5 mM sodium L-ascorbic acid (95210, Merck), 500 nM THPTA (CLK-1010, Jena Bioscience), 100 $\mu$M $CuSO_4$ (7758-98-7, ACROS Organics™) and 100 mM HEPES (pH 7.25) 5h at 37 °C. The total volume of the click reaction was 30 $\mu$L with 26 $\mu$L of cell lysate. To the click mixture 90 $\mu$L methanol, 22.5 $\mu$L chloroform and 60 $\mu$L $dH_2O$ were added with brief vortex steps in between additions. Reaction mixture was centrifuged at 20000 x g for 5 min and the upper aqueous part was discarded without disturbing the interface layer that contains the proteins. Subsequently, 67.5 $\mu$L methanol were added, briefly vortexed and centrifuged at 20000 x g for 5 min. The supernatant was carefully removed and 450 $\mu$L methanol were added, briefly vortexed and centrifuged at 20000 x g for 5 min. Supernatant was removed and protein pellets were air-dried for 30 min. Dried protein pellets were resuspended in SDS buffer containing Orange

G (50 mM Tris (pH6.8), 0.5% SDS, 2.5% glycerol, 1.25% β-Mercaptoethanol, 0.01% Orange G) and afterwards loaded on a NuPAGE 4-12% Bis-Tris gel (NP0322, invitrogen) and run for 90 min at 130 V. The fluorescent standard (LC5928, ThermoFisher Scientific) was diluted 1:1000 in 1X Orange G buffer and was loaded on the gel. Gel was imaged with Typhoon FLA 9500 (GE Healthcare) using Cy2 filter settings. Subsequently, the gel was stained with 1X SyproRed (S6653, ThermoFisher Scientific) for 10 min in 7.5% acetic acid solution and afterwards washed for 10 min in 7.5% acetic acid solution. Following the washing step, the gel was imaged using the Typhoon imager using Cy5 filter settings. The generated images were processed in Fiji: Average background intensity was subtracted from the average fluorescence value per lane and normalised to protein intensity in the same lane after background subtraction. Afterwards the ratios of +UV to -UV were calculated. The experiment was repeated 4 times resulting in 4 gels per condition.

**Image Acquisition and Analysis**

Images were either acquired on a single photon point scanning confocal system (Zeiss LSM 880 Airy inverted) using Zeiss Plan-Apochromat 63x 1.4 Oil DIC objective (all Figures except for figure 4f) or an Olympus IX83 with a Yokogawa W1 CSU with SoRa using an Olympus 100X 1.5 Oil objective. Images were analysed using a custom-made Python script. Below, a brief description of the image analysis workflow is given, the source code including a detailed readme file and all imaging data used for the quantifications are provided in a publicly accessible repository via a link in the main text manuscript.

The set of images containing the DAPI, plasma membrane and sugar stain (Figures 1, 2) were analysed as follows: The nuclear channel was segmented by first applying a Gaussian blur (sigma = 3) and then a Li threshold (skimage package) to generate a binary mask. The holes in the binary mask were filled using the ndimage package. To segment the membrane signal, a Gaussian blur was first applied (sigma = 3). Depending on the cell type, true signal was defined by subtracting the mean signal plus either a quarter (HCT116), half (MDCK) or two (MIN6) standard deviations. To this resulting image, a local threshold was applied with a block size of 15 to generate a binary mask (skimage package). Small objects were removed (minimum size = 250). Regions of the nuclear mask that overlapped the membrane mask were removed from the nuclear mask. Finally, the sugar channel was used to segment the cells. To this purpose the cells were delimited using the membrane mask after applying a gaussian blur (sigma = 5). Cells were then segmented using the cellpose package to generate a labelled cytoplasm mask. The nucleus mask was then labelled with its corresponding cytoplasm. The nuclear and membrane regions were then removed from this mask. Using these masks, the mean intensity for the membrane, cytoplasm and nucleus was calculated for all timepoints, probes and UV conditions using the raw, uncorrected sugar images.

The images displaying nuclear compartments were analysed by first segmenting the nuclear channel. As before, a Gaussian blur (sigma = 3) was applied followed by a Li threshold (skimage package) to generate a binary image of the DAPI signal. The holes in the mask were filled and then the mask was eroded with a 5x5 matrix. To give a unique label to the nuclei we used cellpose on the original nuclear image. This cellpose mask was then multiplied with the eroded binary mask, this was the final nuclear mask. The antibody signal was then segmented using a Gaussian filter (sigma = 3) and an isodata threshold. To preserve the correspondence of the antibody signal to its corresponding nucleus, the antibody mask was multiplied by the nuclear mask to generate a labelled antibody mask. The regions that corresponded to the antibodies were then subtracted from the nuclear mask to generate nuclear masks without the dense fibrillar component, the nuclear speckles or the H3K9me3 positive regions, respectively. Since we observed that irradiation of the DAPI dye with the 488 nm laser line resulted in a low-intensity background signal in the green channel, we corrected for this effect. We used control images of the cells that were not treated with with bifunctional sugar probes to calculate the extent of the DAPI into the green channel. A linear regression of signal in the green channel against signal in the DAPI channel (blue) was fitted (See SI Figure 9 for details). The background signal in the sugar images was removed by subtracting an image generated using the DAPI channel and the calculated linear fit. Finally, the ratio between the mean intensity of the antibody labelled region (nucleoli or nuclear speckles) and the rest of the nucleus was calculated.

**Data presentation and processing statement**

The same dataset (consists of two independent repetitions of the experiments) was used in Figure 1 and 2. All images were acquired with the same settings (using the brightest images fluorescence intensities, which were observed for MIN6 360 min time point as a reference for setting the dynamic range of the acquisition). For visualization purposes, images of HCT116 cells were brightness-contrast adjusted, as the observed intensities for the different cell lines were very different and spatial details are difficult to discern in the low-intensity images. Representative images for all experimental conditions are shown in SI Figures 3-5 for comparison. Care was taken to not oversaturate any pixels by setting the maximum pixel value to the highest observed pixel intensity in the original image. For SI Figures 3.1-5.1, brightness-contrast adjusted images are shown to better illustrate developments in the time course experiments for individual cell lines. SI Figures 3.2-5.1 show unadjusted raw images to illustrate differences in staining intensities between different cell lines. All original images are deposited in a public repository, which is accessible via a link given in the main text. Data analysis was carried out in all cases on unadjusted images. The dataset for Figure 3 (consisting of two independent repetitions of the experiments) was acquired with adjusted acquisition settings for the monosaccharide probe channel compared to the dataset displayed in Figures 1 and 2, as this dataset encompassed only experiments with HCT116 cells. Image processing for visualization was carried out using Fiji. Images for Figure 3 had to be acquired with new settings (change of lasers during the revision processes) and are not directly comparable in fluorescent intensity to images in other figures. Data for Figure 3 consists of two independent repetitions of the experiment. Images for Figure 3 were analyzed with the same python script as used for Figure 1 and 2.

**Chemical Synthesis**

All chemicals were obtained from commercial sources (Acros, Sigma-Aldrich, TCI chemicals, Alfa Aesar, Roth, Fluka or Merck) and were used without further purification. Solvents for flash chromatography were obtained from VWR and dry solvents were obtained from Sigma. Deuterated solvents were obtained from Deutero GmbH, Karlsruhe, Germany. TLC was performed on precoated plates of silica gel (Merck, 60 F254) using UV light (254 or 365 nm) or a solution of phosphomolybdic acid in EtOH (10 g phosphomolybdic acid, in 100 mL EtOH) for analysis. Preparative column chromatography was performed using silica gel from Merck, Darmstadt, Germany (silica 60, grain size 0.063-0.200 mm) with a pressure of 1.5 bar. Detailed purification conditions are given for the respective compounds. $^{1}$H- and $^{13}$C-NMR-spectra were measured on 400 MHz Advance™ III HD Nanobay Bruker spectrometer. Chemical shifts of $^{1}$H- and $^{13}$C-NMR-spectra are referenced indirectly to tetramethylsilane. *J* values are given in Hz and chemical shifts in ppm. Splitting patterns are designated as follows: s, singlet; d, doublet; t, triplet; q, quartet; m, multiplet; b, broad. $^{13}$C-NMR-spectra were broadband hydrogen decoupled. Mass spectra were recorded using a QExactive Orbitrap FTMS instrument (Thermo Fisher Scientific) equipped with a robotic nanoflow electrospray ion source. FTMS were acquired for the mass range of m/z 200-1500 with the target mass resolution set to 140000 at m/z 200. Samples were dissolved in a solution of $^{i}$PrOH/MeOH/$CHCl_3$ 4:2:1 containing 7.5 mM ammonium formiate. The spectra were evaluated using the Xcalibur Qual Browser software.

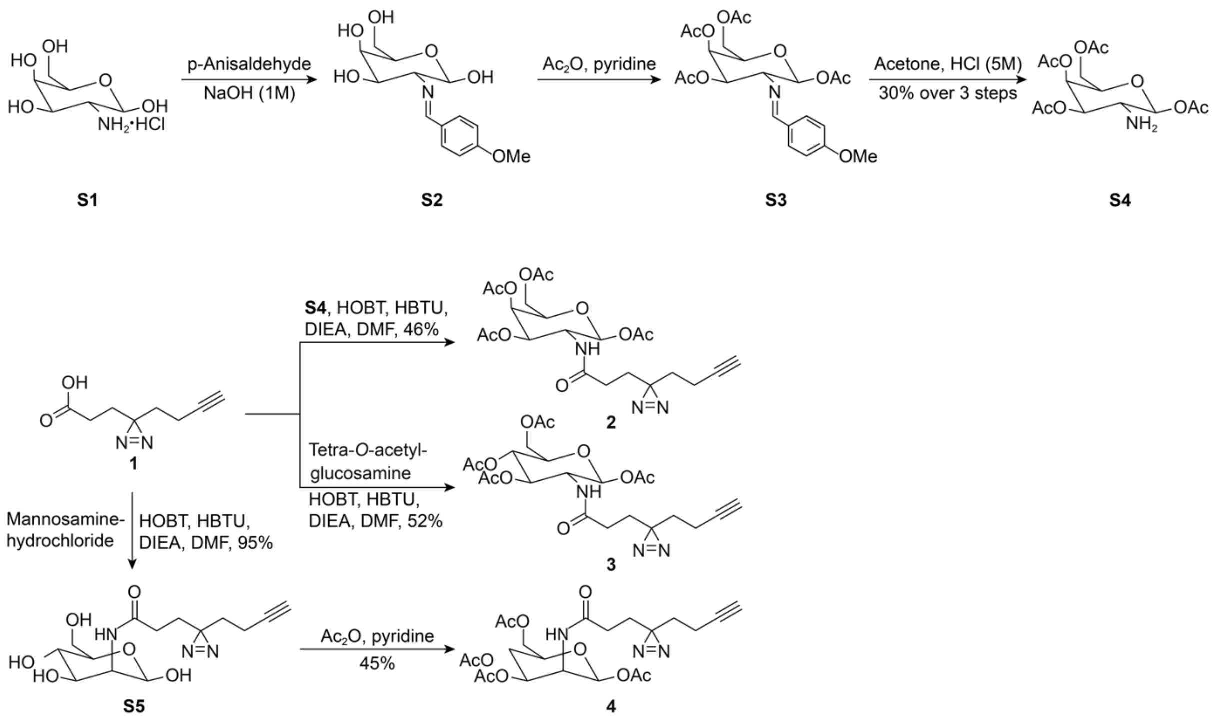

**SI Scheme 1 |** Synthesis of probes **2**,**3 and 4**.

The minimal bifunctional linker **1** was synthesized according to a previously reported protocol[2]. The synthesis of 1,3,4,6,-*O*-Tetraacetylgalactosamine was carried out by slightly adapting a procedure for the synthesis of Tetra-*O*-acetylglucosamine reported by Biswas et al[3].

2-*N*-Methoxybenzylidene-galactosamine (**S2**)

A chilled (0 °C) solution of galactosamine hydrochloride (**S1**) (500 mg, 2.32 mmol) in NaOH (2.5 ml, 1M) was treated with *p*-anisaldehyde (0.338 ml, 2.41 mmol) and subsequently stirred for 1 h. The resulting colourless precipitate was collected by filtration and subsequently washed with cold water, ethanol and diethyl ether (1.5 ml each). The title compound was obtained as a colourless solid and used immediately without further purification.

2-*N*-Methoxybenzylidene-1,3,4,6,-O-tetraacetyl-galactosamine (**S3**)

A solution of **S2** (550 mg, 1.85 mmol) in pyridine (4 ml) was stirred for 5 min at 0 °C under inert conditions and subsequently treated with 0.945 ml acetic anhydride (10.0 mmol). The reaction mixture was stirred at 0 °C for 2 h and allowed to reach room temperature overnight. After addition of water (20 ml), the resulting precipitate was collected by filtration, washed with cold water (3 x 1 ml) and dried under vacuum. The title compound was obtained as a colourless solid and used immediately without further purification.

1,3,4,6,-O-Tetraacetyl-galactosamine (**S4**)

A solution of **S3** (600 mg, 129 mmol) in acetone (5 ml) was treated with 5M HCl (0.3 ml). The reaction mixture was stirred for 30 min and subsequently treated with cold diethyl ether (25 ml) and stirring was continued for an additional hour at 0 °C. The resulting precipitate was collected by filtration and washed with cold diethyl ether. The crude material was purified using flash column chromatography on silica using the eluent $CHCl_3/MeOH/H_2O$ 65:35:1. The product was obtained as a colourless solid in a yield of 250 mg (72 mmol, 30% over three steps).

$^1$H NMR (400 MHz, DMSO-d6): δ = 8.68 (s, 2H), 5.89 (d, *J* = 8.7 Hz, 1H), 5.31 – 5.23 (m, 2H), 4.31 (dd, *J* = 6.2, 6.2 Hz, 1H), 4.11 – 3.98 (m, 2H), 3.45 – 3.38 (m, 1H), 2.17 (s, 3H), 2.13 (s, 3H), 2.01 (s, 3H), 2.00 (s, 3H) ppm.
$^{13}$C NMR (101 MHz, DMSO-d6): δ = 170.38, 170.35, 169.79, 169.11, 90.73, 71.52, 69.29, 66.23, 61.63, 49.79, 21.29, 21.14, 20.97, 20.81ppm.
HR-MS (ESI positive) m/z calculated for $C_{14}H_{21}NO_9$: 347.12; found: 348.127 $[M+H]^+$.

2-*N*-(3'*H*-Diazirine)-7'-octynyl-mannosamine (**S5**)

22.5 mg of **1** (135.4 µmol, 1.1 eq.) were dissolved in 2 ml dry DMF. 81.1 mg HBTU (213.8 µmol,1.8 eq.), 6.3 mg HOBT (46.6 µmol, 0.4 eq.) and 0.05 ml DIPEA (286.3 µmol, 2.4 eq.) were added and the fatty acid was activated for 10 minutes. 25.5 mg of Mannosamine hydrochloride (118.3 µmol, 1 eq.) were added and the reaction mixture was stirred for 3 h at room temperature. Subsequently, DMF was removed under reduced pressure and the crude product was purified by flash chromatography (eluents 10% MeOH/DCM). The product was isolated as a white solid in a yield of 95% (37.0 mg, 113.0 µmol). The product is a mixture of the α and β isomer (ratio around 3:7).

$^{1}$H NMR (400 MHz, MeOD) δ 4.41 – 4.36 (m, 0.3H), 4.32 – 4.23 (m, 0.7H), 3.99 (dd, *J* = 9.7, 4.7 Hz, 0.7H), 3.89 – 3.69 (m, 3H), 3.66 – 3.52 (m, 1H), 3.53 – 3.41 (m, 0.5H), 3.28 – 3.17 (m, 0.7H), 2.82 (s, 1H), 2.27 (d, *J* = 2.6 Hz, 1H), 2.22 – 2.08 (m, 2H), 2.08 – 1.94 (m, 2H), 1.76 (t, *J* = 7.8 Hz, 2H), 1.69 – 1.56 (m, 2H), 1.41 – 1.27 (m, 4H).*

*The 4 protons attached to the pyranose show different chemical shifts for the α and β isomer

$^{13}$C NMR (101 MHz, MeOD) δ = 176.21, 175.11, 94.95, 94.88, 83.62, 78.20, 74.45, 73.39, 70.59, 70.34, 70.32, 68.43, 68.04, 62.17, 61.99, 55.81, 55.74, 55.00, 43.79, 38.88, 33.27, 33.24, 31.06, 30.93, 29.97, 29.83, 29.00, 28.95, 19.29, 13.85, 13.81, 13.17 ppm.*

*Some carbons of the α and β isomer give individual signals.

HR-MS (ESI positive) m/z calculated for $C_{14}H_{21}N_3O_6$: 327.1430; found: 328.150 [M+H]$^+$.

2-*N*-(3'*H*-Diazirine)-7'-octynyl-1,3,4,6,-*O*-tetraacetyl-galactosamine (**2**)

A flask charged with 5 mg HOBT (37 µmol) and 60 mg HBTU (158 µmol) was treated with a solution of 25 mg **1** (150 µmol) and 100 µl DIEA (611 µmol) in dry DMF (2 ml) and the reaction mixture stirred for 10 min under inert conditions. Subsequently, a solution of **S4** (50 mg, 144 µmol) in 1 ml dry DMF was added and stirring continued overnight. The solvent was removed under reduced pressure and the residue purified by repeated flash column chromatography on silica using the eluent EtOAc followed by $CH_2Cl_2$/MeOH 9:1 for the first column and $CH_2Cl_2$/MeOH 9:1 for the second column. The title compound was isolated as a colourless solid in a yield of 34 mg (69 µmol, 48 %).

$^{1}$H NMR (400 MHz, $CDCl_3$): δ = 5.66 (d, *J* = 8.8 Hz, 1H), 5.42 (d, *J* = 9.4 Hz, 1H), 5.31 (d, *J* = 1.1 Hz, 1H), 5.04 (dd, *J* = 11.3, 3.3 Hz, 1H), 4.41 – 4.29 (m, 1H), 4.16 – 4.02 (m, 2H), 4.02 – 3.94 (m, 1H), 2.11 (s, 3H), 2.08 (s, 3H), 2.00 – 1.90 (m, 9H), 1.85 – 1.72 (m, 4H), 1.55 (t, *J* = 7.2 Hz, 2H) ppm.

$^{13}$C NMR (101 MHz, $CDCl_3$): δ = 171.29, 170.78, 170.44, 170.19, 169.61, 92.84, 82.72, 71.86, 70.26, 69.36, 66.38, 61.32, 49.81, 32.56, 30.20, 27.86, 27.55, 20.97, 20.69 (multiple overlapping peaks), 13.24ppm.

HR-MS (ESI negative) m/z calculated for $C_{22}H_{29}N_3O_{10}$: 495.19; found: 540.184 [M+HCOO]$^-$.

2-*N*-(3'*H*-Diazirine)-7'-octynyl-1,3,4,6,-*O*-tetraacetyl-glucosamine (**3**)

A flask charged with 5 mg HOBT (37 µmol) and 60 mg HBTU (158 µmol) was treated with a solution of 25 mg **1** (150 µmol) and 100 µl DIEA (611 µmol) in dry DMF (2 ml) and the reaction mixture stirred for 10 min under inert conditions. Subsequently, a solution of 1,3,4,6,-*O*-tetraacetyl-glucosamine (50 mg, 144 µmol) in 1 ml dry DMF was added and stirring continued overnight. The solvent was removed under reduced pressure and the residue purified by repeated flash column chromatography on silica using the eluent EtOAc followed by $CH_2Cl_2$/MeOH 9:1 for the first column and $CH_2Cl_2$/MeOH 9:1 for the second column. The title compound was isolated as a colourless solid in a yield of 37 mg (75 µmol, 52 %).

$^{1}$H NMR (400 MHz, $CDCl_3$): δ = 5.73 (d, *J* = 8.7 Hz, 1H), 5.47 (d, *J* = 9.3 Hz, 1H), 5.21 – 5.11 (m, 2H), 4.34 – 4.23 (m, 2H), 4.15 (dd, *J* = 12.5, 2.2 Hz, 1H), 3.86 – 3.78 (m, 1H), 2.21 – 2.00 (m, 15H), 1.94 – 1.79 (m, 4H), 1.64 (t, *J* = 7.0 Hz, 2H) ppm.

$^{13}$C NMR (101 MHz, $CDCl_3$): δ = $^{13}$C NMR (101 MHz, $CDCl_3$) δ 171.25, 171.04, 170.68, 169.56, 169.26, 92.55, 82.71, 72.96, 72.42, 69.35, 67.66, 61.62, 53.07, 32.56, 30.13, 27.81, 27.54, 20.96, 20.75, 20.71, 20.60, 13.25 ppm.

HR-MS (ESI negative) m/z calculated for $C_{22}H_{29}N_3O_{10}$: 495.19; found: 540.185 [M+HCOO]$^-$.

2-*N*-(3'*H*-Diazirine)-7'-octynyl-1,3,4,6-*O*-tetraacetyl-mannosamine (**4**)

18.0 mg of **S5** (55.0 µmol, 1 eq.) was dissolved in 2 ml dry pyridine. 0.05 ml acetic anhydride (53.9 µmol, 1 eq.) were added and the reaction mixture was stirred overnight at room temperature. The next morning, additional 0.05ml $Ac_2O$ (53.9 µmol, 1 eq.) were added and the mixture was stirred for more 2 h. All volatiles were removed under reduced pressure and the crude product was purified by flash chromatography (eluents 1% MeOH/chloroform). The product was isolated as a colorless oil in a yield of 48 % (13.1 mg, 26.4 µmol) as a mixture of the α and β isomer (ratio around 1:1).

$^1$H NMR (400 MHz, $CDCl_3$) δ= 6.03 (s, 0.5H), 5.85 (s, 0.5H), 5.83 – 5.73 (m, 1H), 5.33 (dd, *J* = 10.2, 4.4 Hz, 0.5H), 5.23 – 5.09 (m, 1H), 5.04 (dd, *J* = 9.9, 4.0 Hz, 0.5H), 4.81 – 4.71 (m, 0.5H), 4.67 – 4.56 (m, 0.5H), 4.35 – 4.24 (m, 1H), 4.14 – 3.99 (m, 1.5H), 3.85 – 3.74 (m, 0.5H), 2.17 (s, 2H), 2.11 (s, 5H), 2.09 – 1.96 (m, 10H), 1.96 – 1.83 (m, 2H), 1.66 (t, *J* = 7.0 Hz, 2H) ppm.*

*The 4 protons attached to the pyranose show different chemical shifts for the α and β isomer

$^{13}$C NMR (101 MHz, $CDCl_3$) δ = 171.96, 171.48, 170.71, 170.23, 170.15, 169.78, 168.48, 168.27, 91.74, 90.76, 82.93, 82.89, 73.62, 71.50, 70.28, 69.55, 69.50, 69.00, 65.48, 65.27, 62.08, 61.95, 49.72, 49.53, 32.57, 32.54, 30.50, 30.42, 28.45, 28.42, 27.93, 21.01, 20.91, 20.86, 20.80, 13.43 ppm.*

*Some carbons of the α and β isomer give individual signals.

HR-MS (ESI negative) m/z calculated for $C_{22}H_{29}N_3O_{10}$: 495.1853; found: 540.185 $[M+HCOO]^-$.

#### Supporting Information – Figures

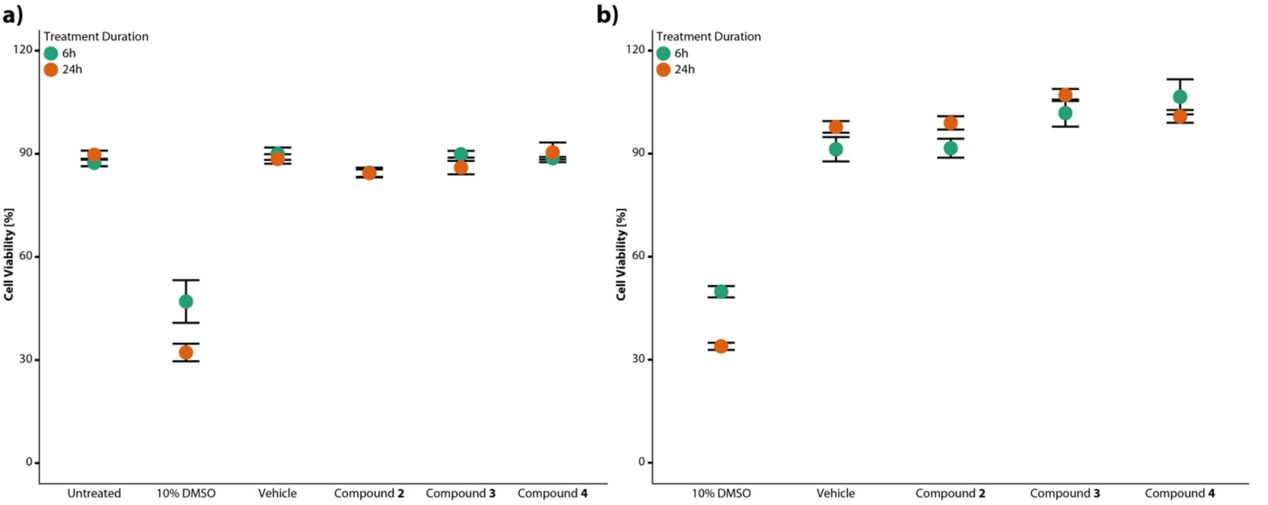


**SI Figure 1 |** Cells remain viable after treatment with the sugar probes. **a**) Cell viability measurement by trypan blue. Cells were treated with the respective compound and control compounds for 6h or 24h and afterwards the percentage of living cells was determined using Trypan blue staining, n=3. **b**) Metabolic activity/cell viability of the cells was assessed using Resazurin. Cells were treated with the respective compound and control compounds for 6h or 24h and afterwards incubated with resazurin assay buffer solution for 1h at 37 °C. Afterwards the fluorescence intensity was measured and the percentage of living cells was calculated with respect to the untreated cells, n = 9. For both experiments, medium containing 10% DMSO was used as a positive control to induce cell death and disturb metabolic activity.

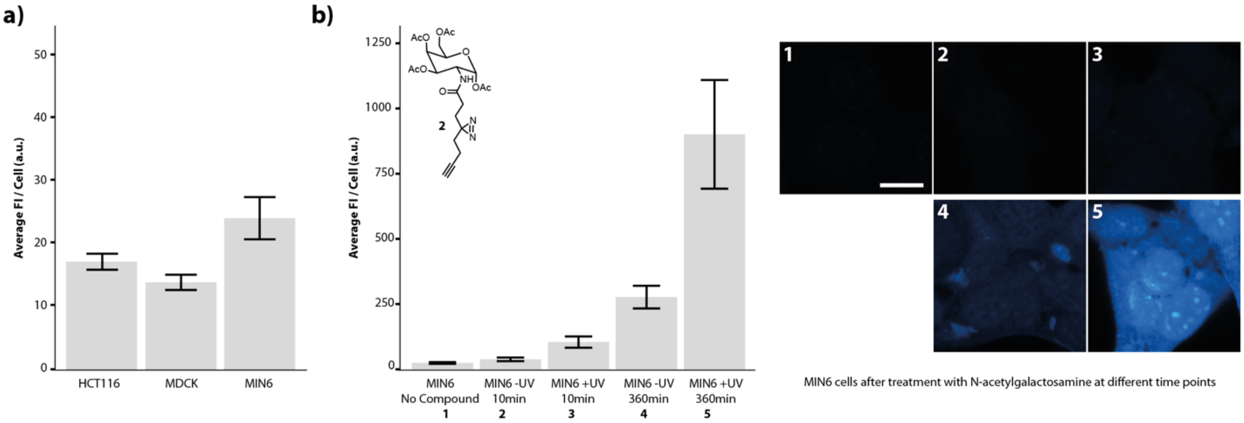


**SI Figure 2 |** Click labelling reactions produce low signal in the absence of probe 2. **a**) Background fluorescence intensities of unspecific click chemistry staining in individual cell lines. Cells were subjected to click chemistry treatments without prior addition of the bifunctional probes. The images were acquired with the same settings as used for data acquisition in Figures 1 and 2. Error bars indicate standard deviation. **b**) Comparison of background fluorescence intensities with observed intensities in MIN6 cells treated with **2** for different time period and under + and -UV conditions. Scale bar = 14 µm.

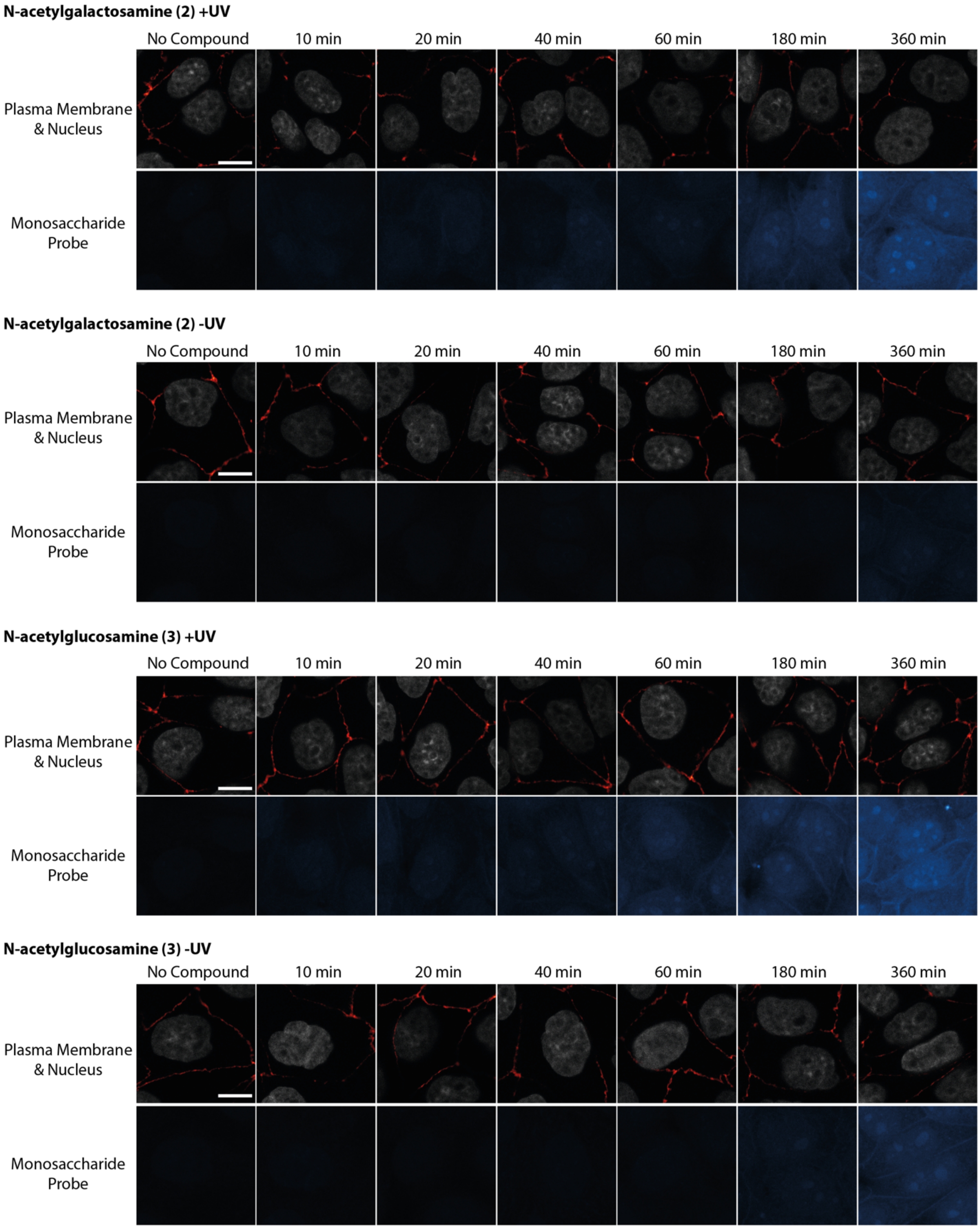


**SI Figure 3-1 |** Monosaccharide uptake of MDCK cells**.** Cells were treated according to the protocol outlined above. Images were acquired with same acquisition settings for all cell lines and brightness-contrast adjusted for better visualization. Unadjusted images of the same cells are shown below in Figure S3-2. Scale bar = 14 µm.

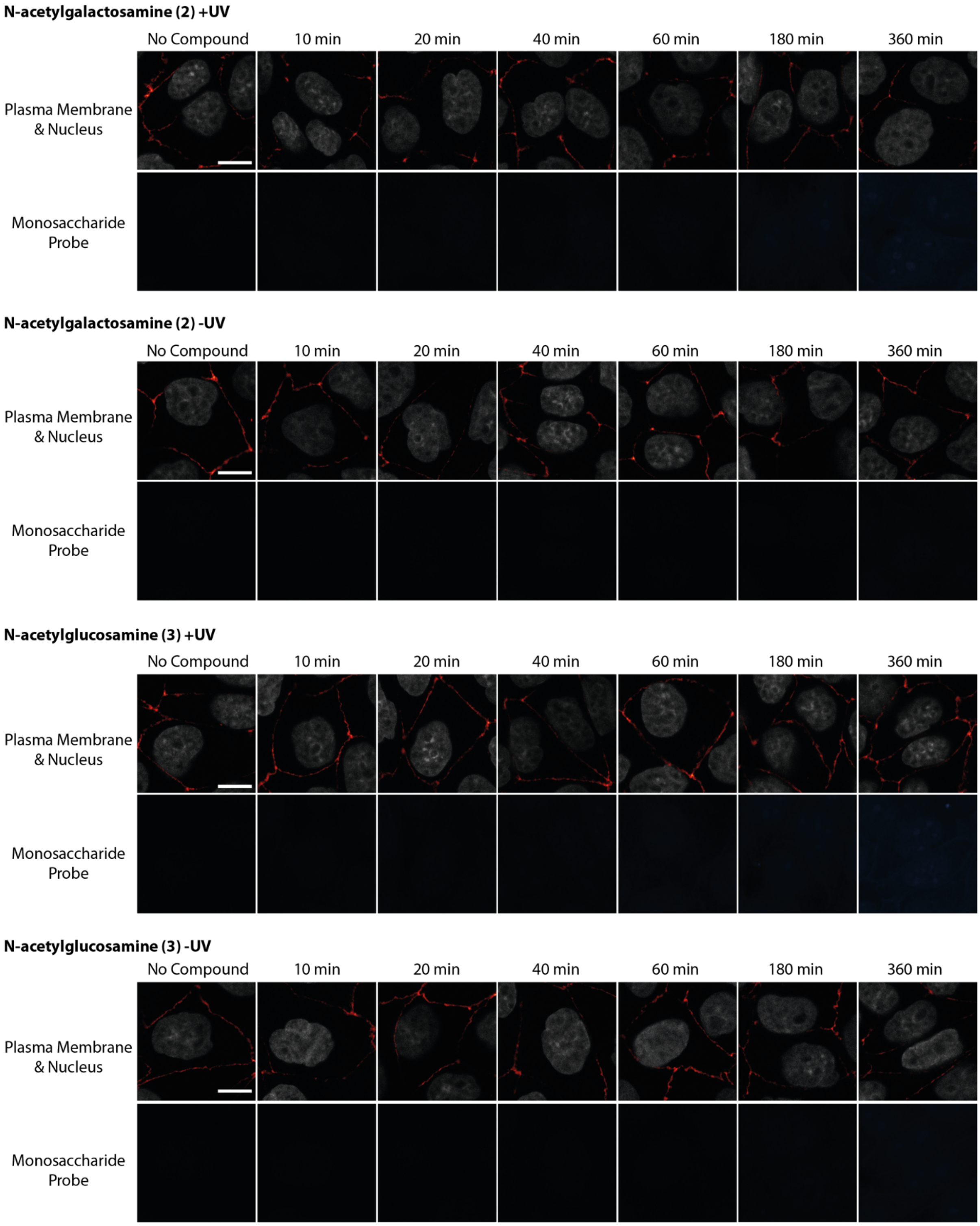


**SI Figure 3-2 |** Monosaccharide uptake of MDCK cells**.** Cells were treated according to the protocol outlined above. Images were acquired with same acquisition settings for all cell lines. Unadjusted images are shown. Brightness-contrast adjusted images of the same cells are shown above in Figure S3-1. Scale bar = 14 µm.

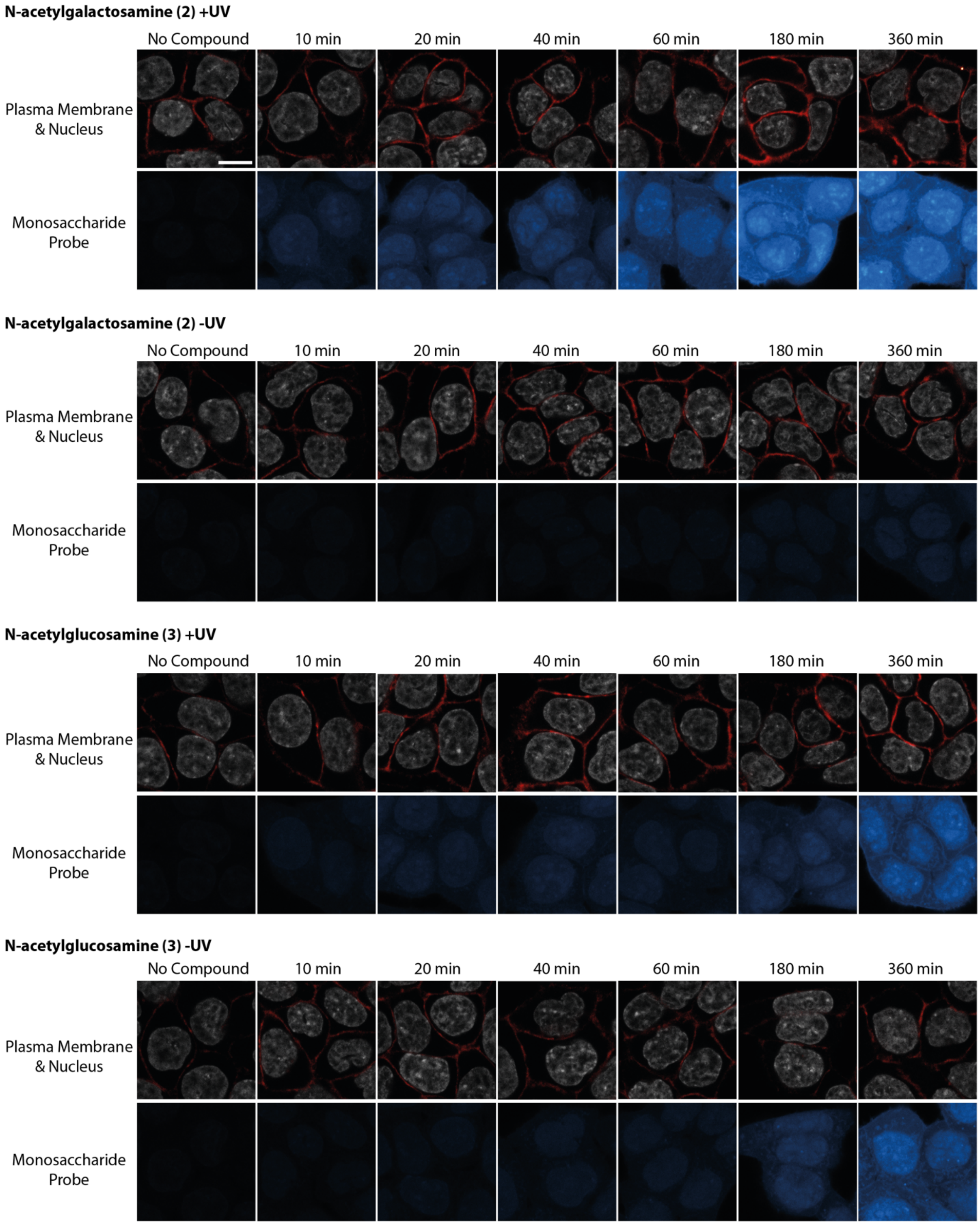


**SI Figure 4-1 |** Monosaccharide uptake of HCT116 cells**.** Cells were treated according to the protocol outlined above. Images were acquired with same acquisition settings for all cell lines and brightness-contrast adjusted for better visualization. Unadjusted images of the same cells are shown below in Figure S4-2. Scale bar = 14 µm.

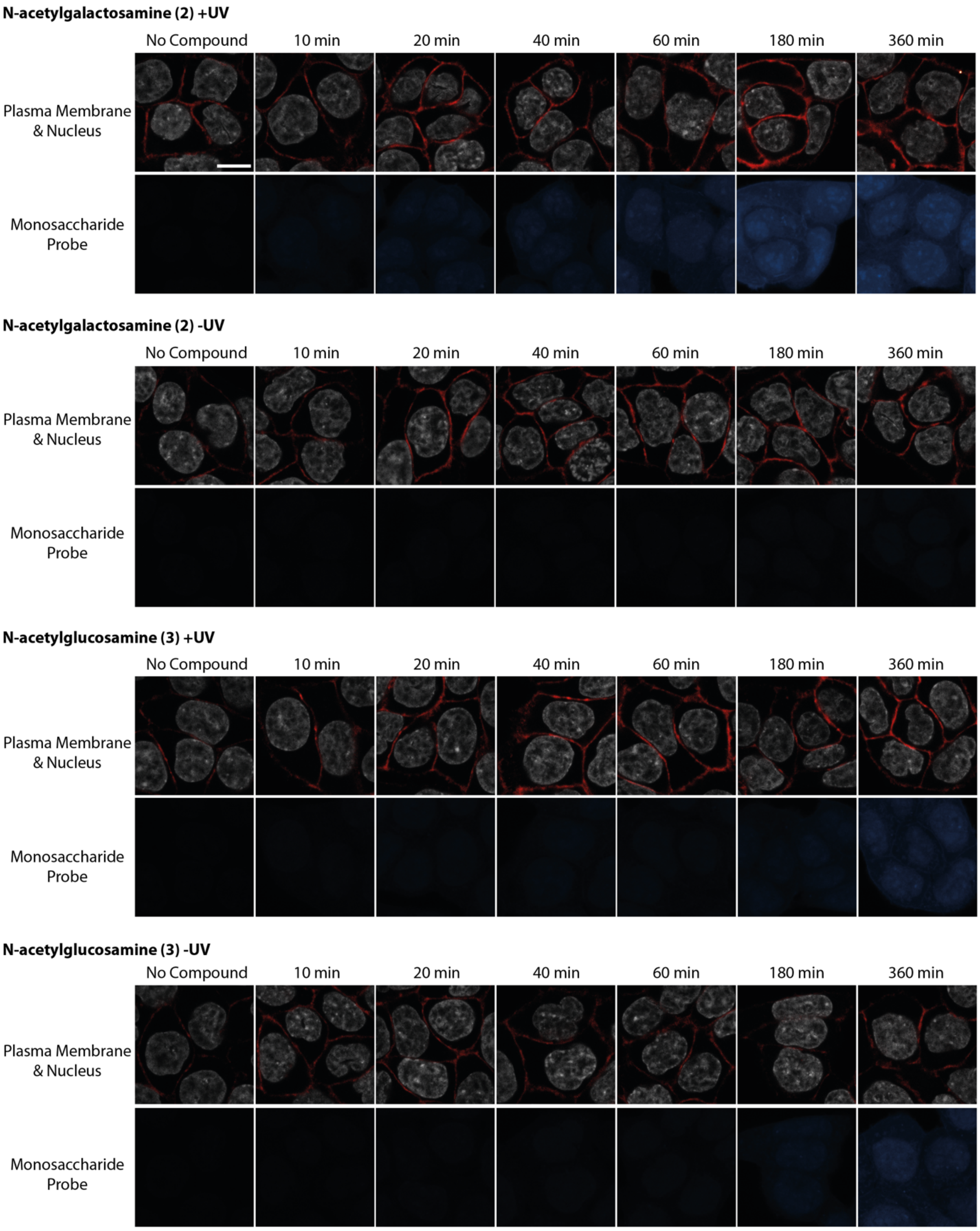


**SI Figure 4-2 |** Monosaccharide uptake of HCT116 cells**.** Cells were treated according to the protocol outlined above. Images were acquired with same acquisition settings for all cell lines. Unadjusted images are shown. Brightness-contrast adjusted images of the same cells are shown above in Figure S4-1. Scale bar = 14 µm.

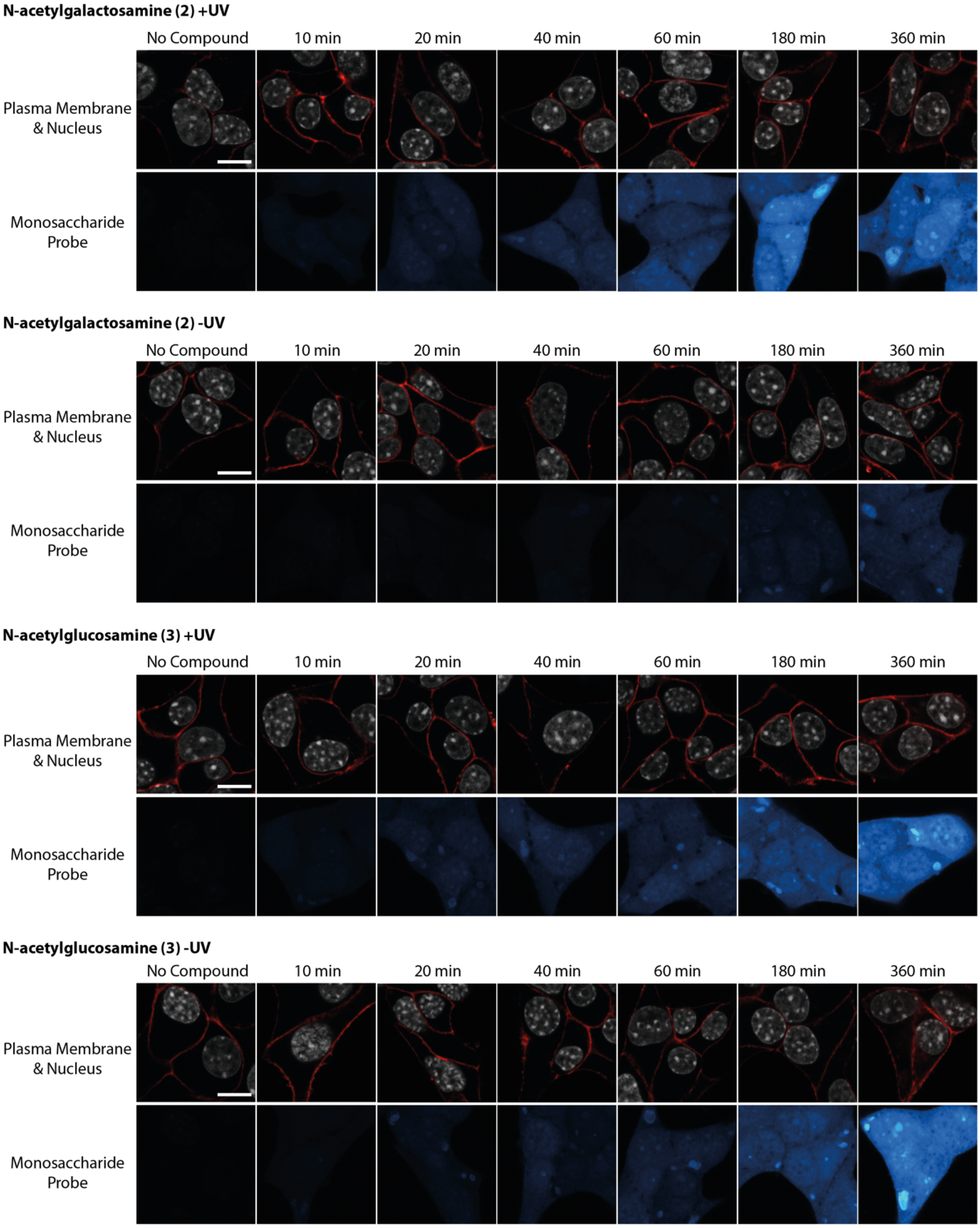


**SI Figure 5-1 |** Monosaccharide uptake of MIN6 cells**.** Cells were treated according to the protocol outlined above. Images were acquired with same acquisition settings for all cell lines. Unadjusted images are shown. Scale bar = 14 µm.

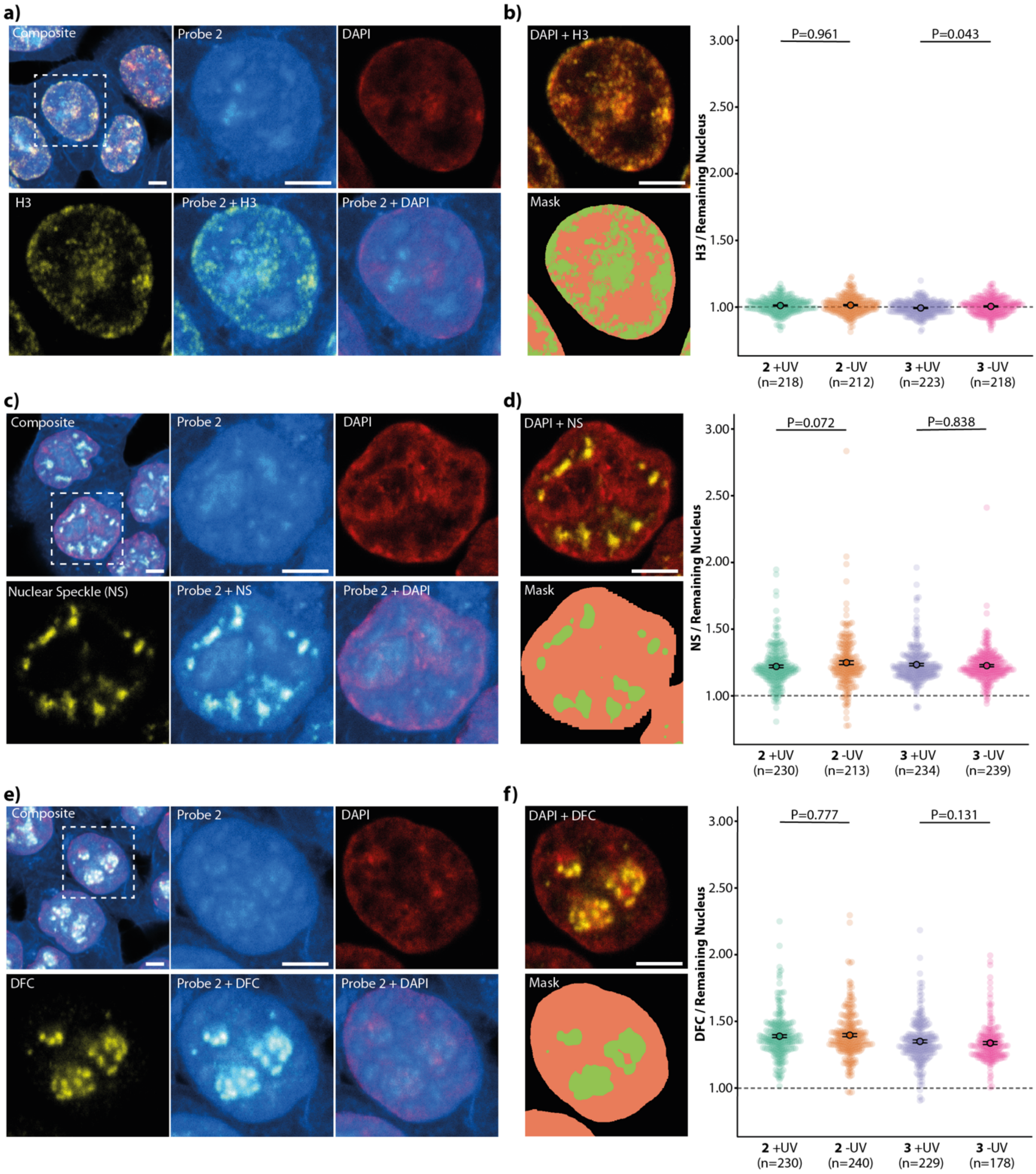


**SI Figure 6 |** Accumulation in nuclear compartments does not differ significantly between + and – UV conditions. Images show the same cells displayed in Figure 3 with additional overlays. Subcellular localisations were analysed with an experiment specific Python macro (see details above). Error bars indicate standard error. Statistical analysis was performed using a two-sided Mann-Whitney-U test. n numbers represent the number of analysed nuclei per condition. Images were acquired with same acquisition settings and brightness-contrast adjusted for better visualization. Scale bar = 10 µm.

6h Pulse Gels

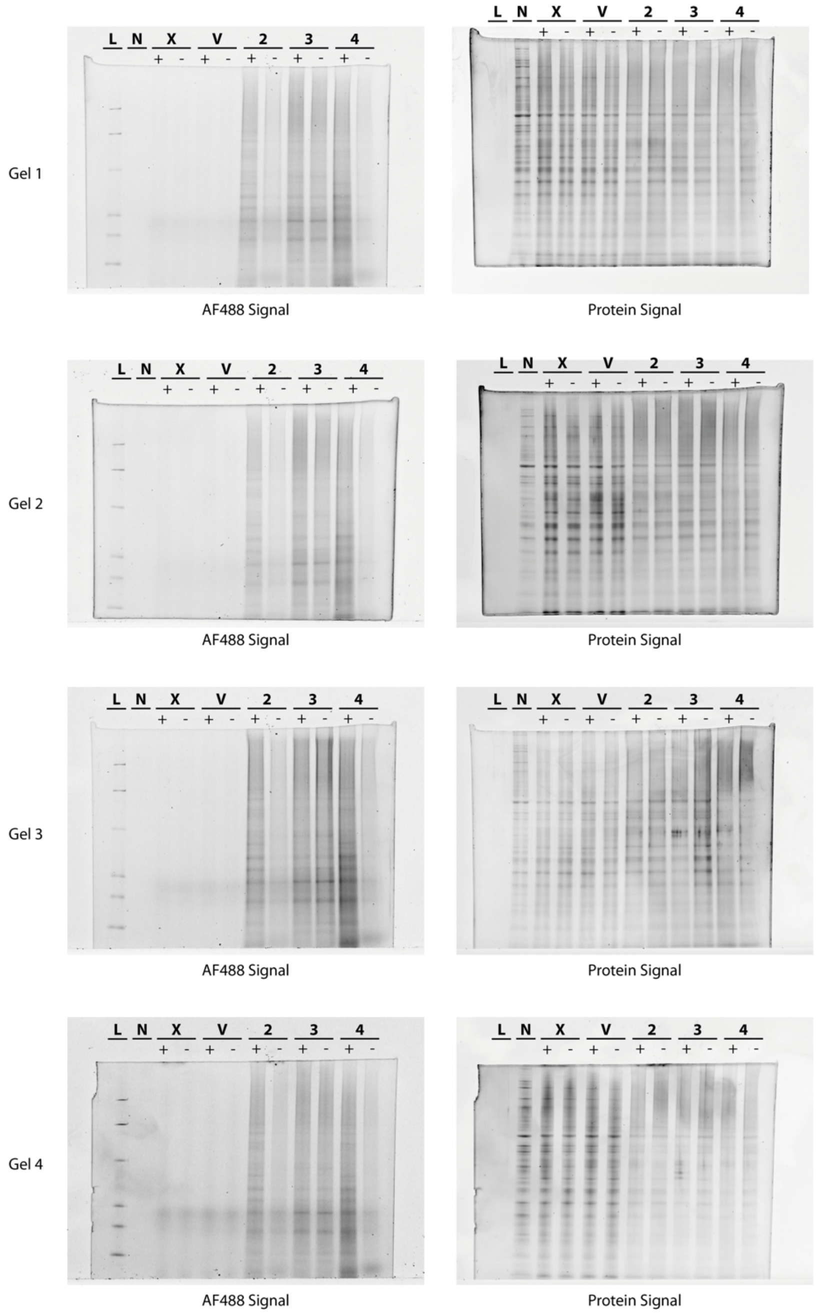


**SI Figure 7 |** Uncut SDS-Page gels of 6h pulse experiment. Cells were treated with 50 µM of probe **2**, **3** or **4** for 6h and afterwards irradiated with 300nm UV light and lysed (+UV) or lysed without irradiation (-UV). The whole cell lysate was subjected to a click labelling reaction and the proteins were analysed by SDS-Page (two-colour gel image). L = Ladder, N = No-click control, X = Untreated and V = Vehicle control. Gel 1 was used in Figure 3. Gels were brightness-contrast adjusted for better visualization.

24h Pulse Gels

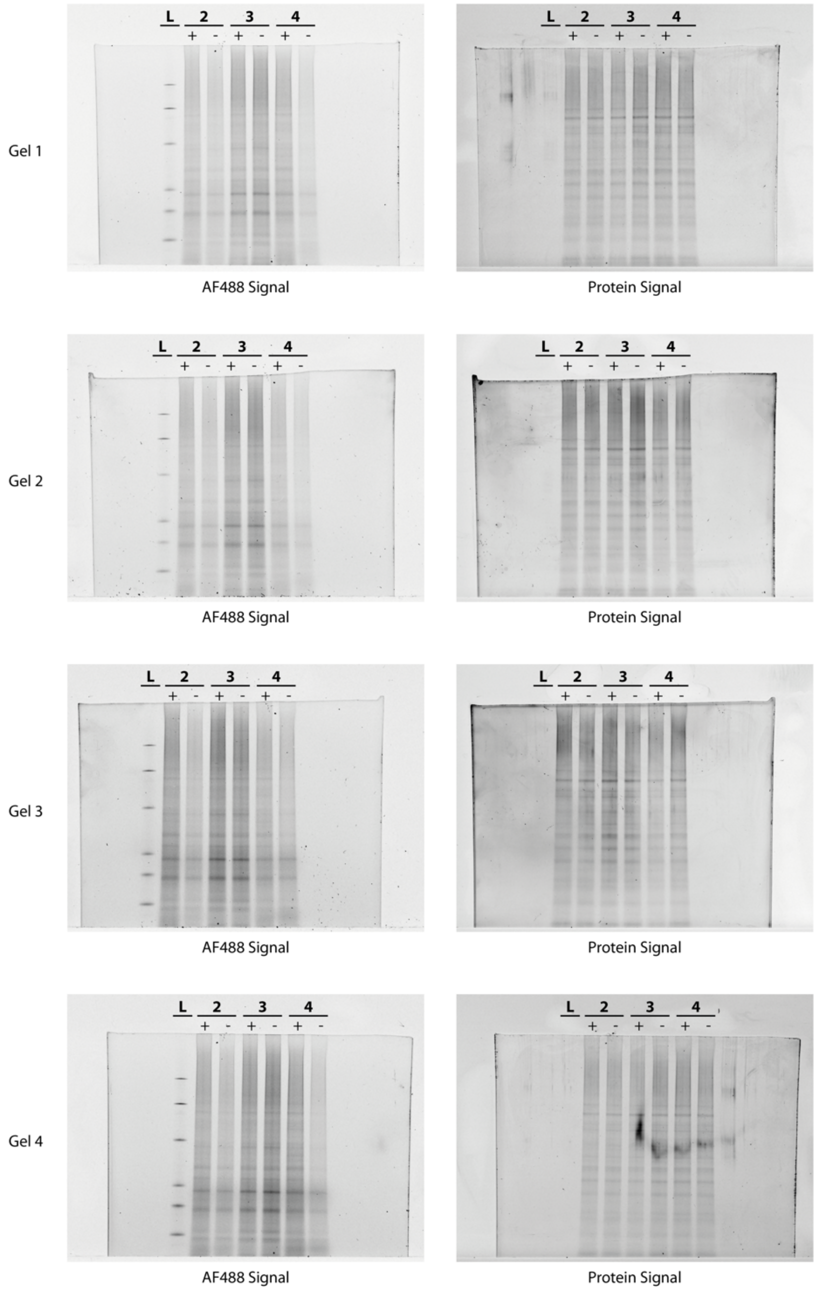


**SI Figure 8 |** Uncut SDS-Page gels of 24h pulse experiment. Cells were treated with 50 µM of probe **2**, **3** or **4** for 24h and afterwards irradiated with 300nm UV light and lysed (+UV) or lysed without irradiation (-UV). The whole cell lysate was subjected to a click labelling reaction and proteins were analysed by SDS-Page (two-colour gel image). L = Ladder, N = No-click control, X = Untreated and V = Vehicle control. Gel 1 was used in Figure 3. Gels were brightness-contrast adjusted for better visualization.

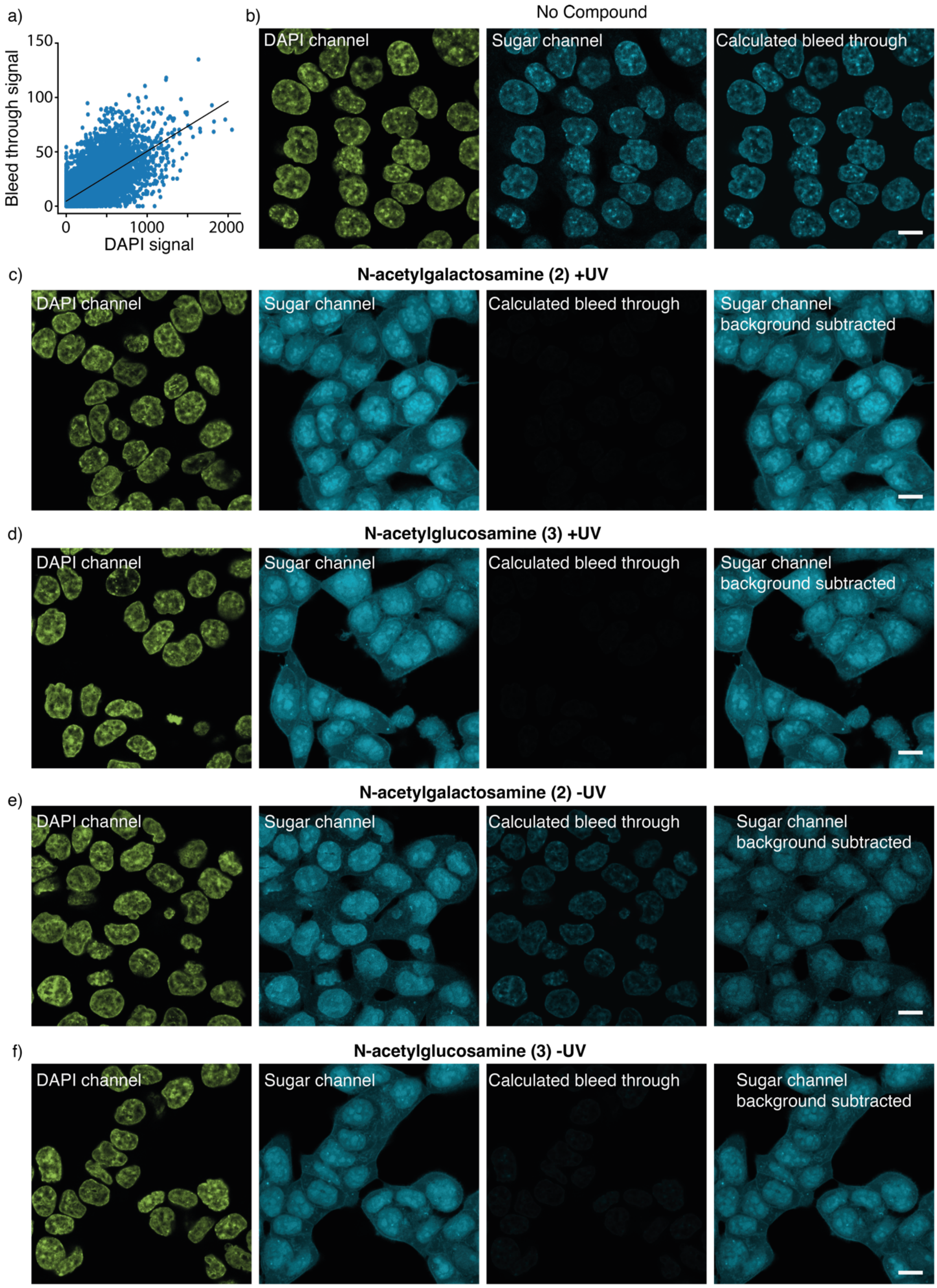


**SI Figure 9 |** Background subtraction for antibody labelled images. **a)** Bleed through signal in the sugar channel from the DAPI staining. To remove the background the amount of bleed through was fitted to a line in order to generate an equivalent bleed through image that was used to subtract the background. **b)** A bleed through image was generated using the linear regression from **a)**, the sugar channel and the calculated bleed through images are shown using the same contrast, the brightness of the DAPI image was adjusted independently. The calculated image adequately represents the amount of bleed through in the no compound condition **c-f)** Using the linear regression the amount of bleed through from the DAPI staining into the sugar channel was estimated for both sugars using the linear fit in **a)**. The fit was used to make a bleed through image from the DAPI image. This image was then subtracted from the sugar channel. Final calculations were carried in the background subtracted image. Scale bar = 10 µm

### Supporting Information – NMR spectra

Compound **S4** $^{1}H$ spectrum

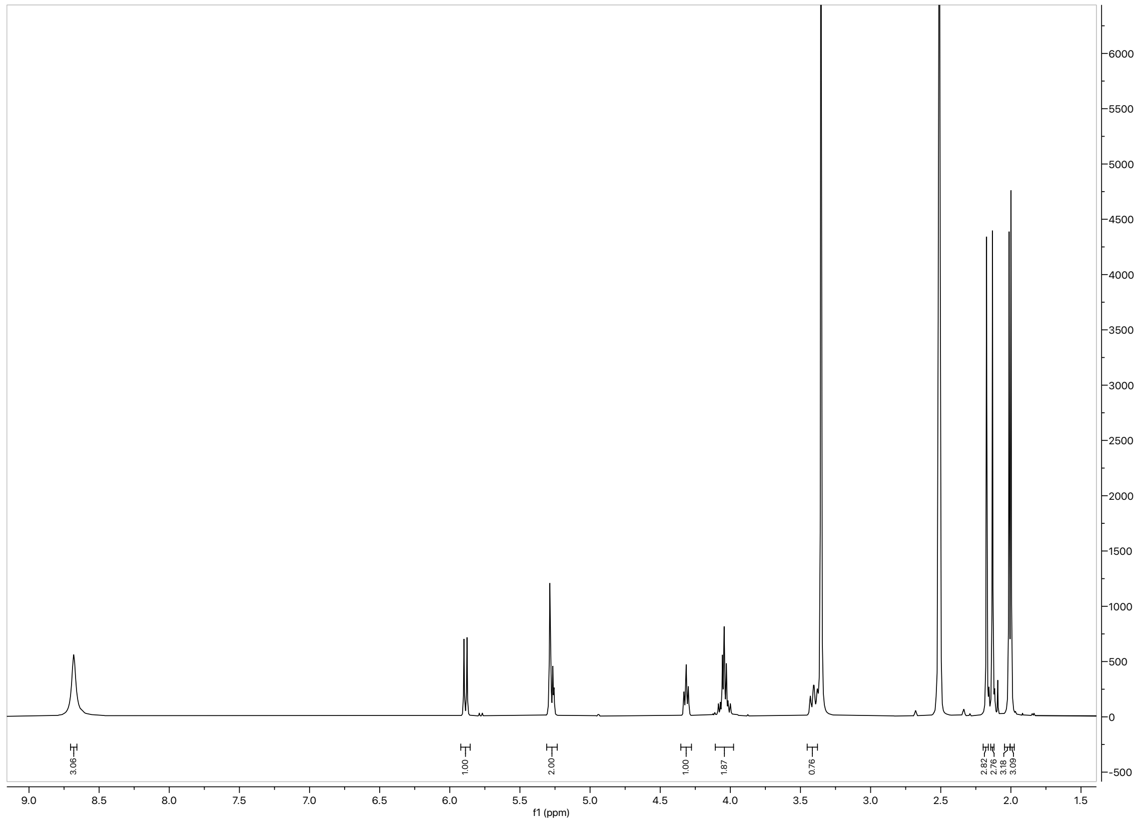

Compound **S4** $^{13}C$ spectrum

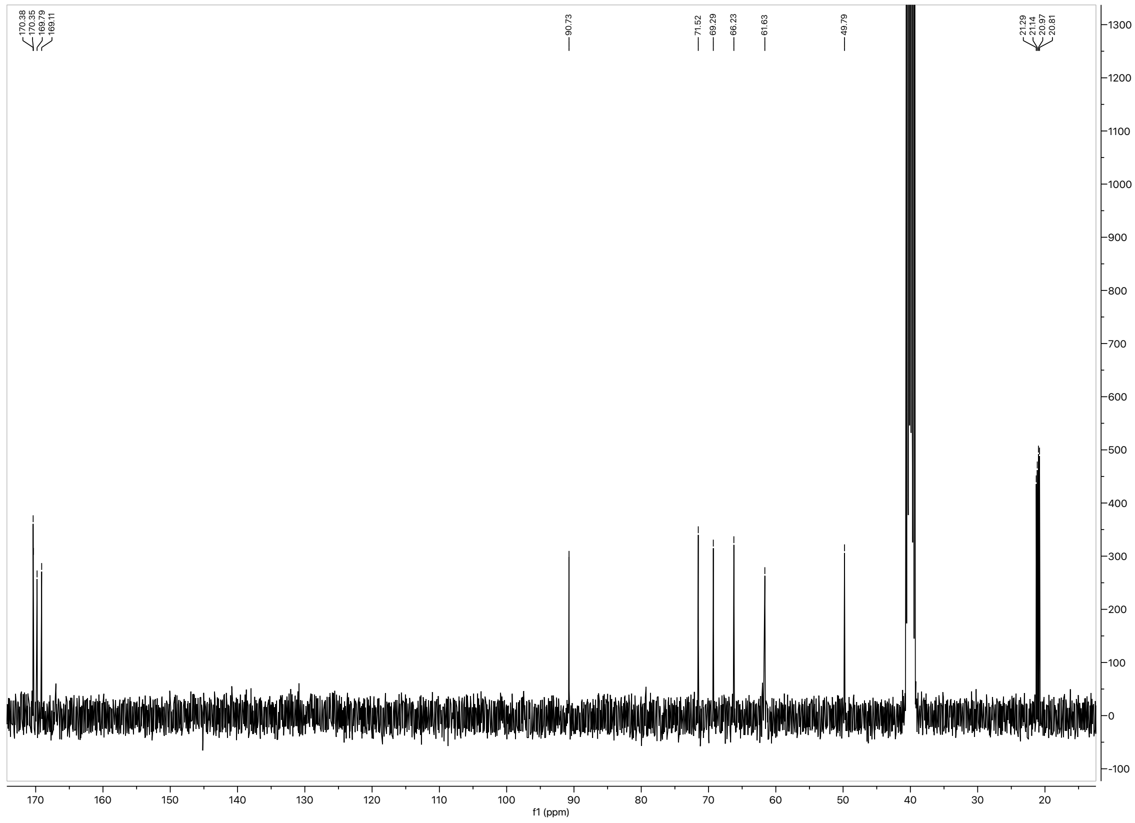

Compound **2** $^{1}$H spectrum

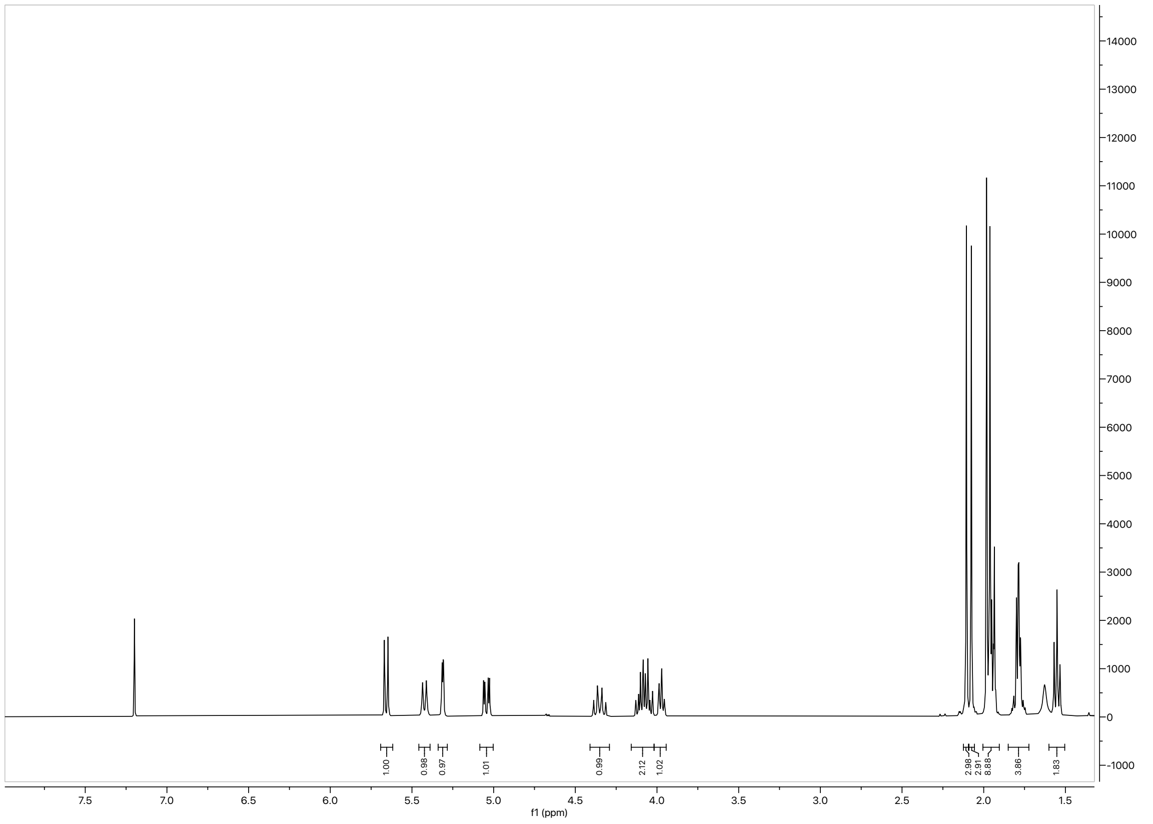

Compound **2** $^{13}$C spectrum

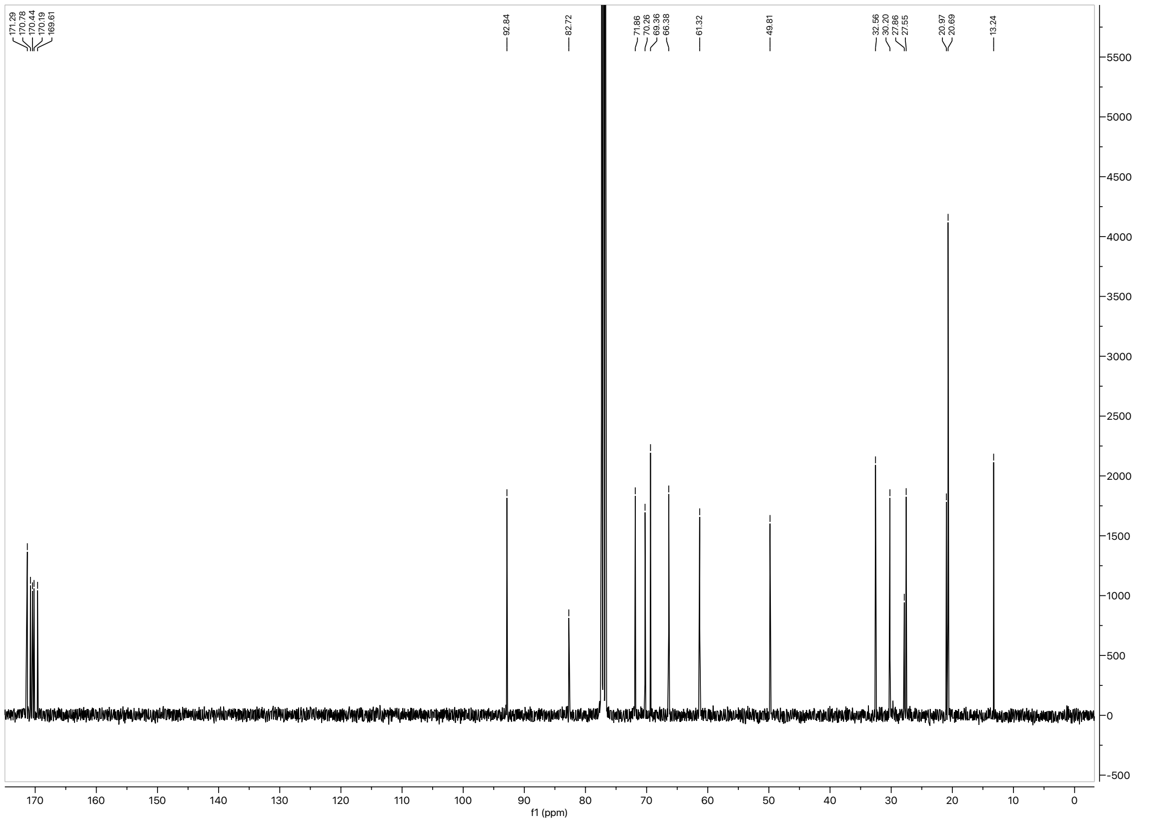

Compound **3** $^{1}$H spectrum

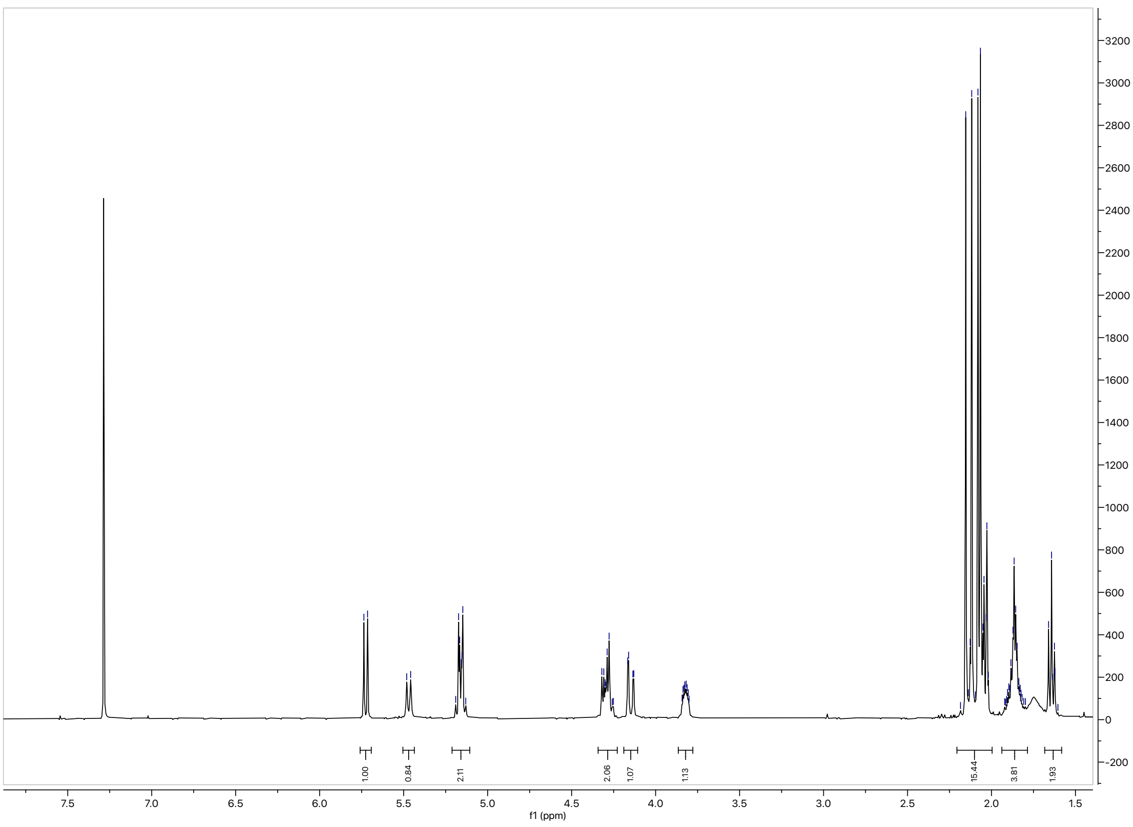


Compound **3** $^{13}$C spectrum

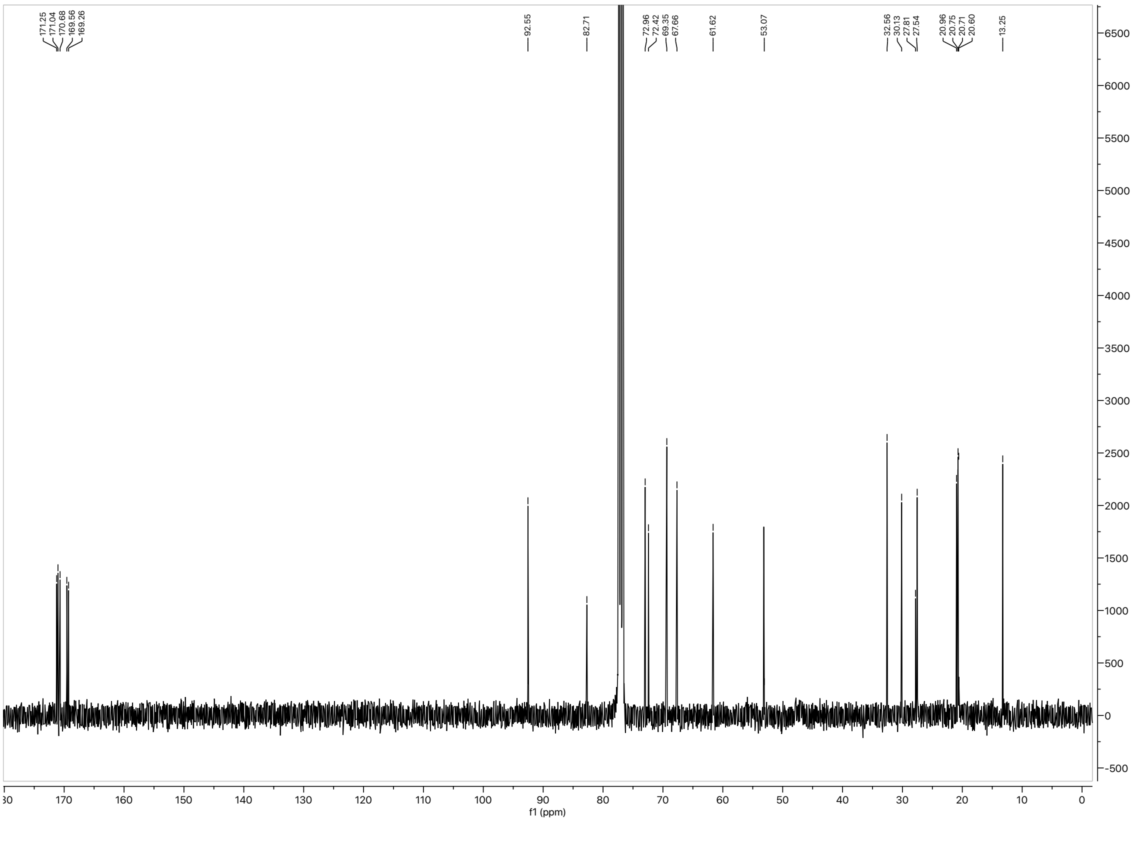

Compound **S5** $^{1}H$ spectrum

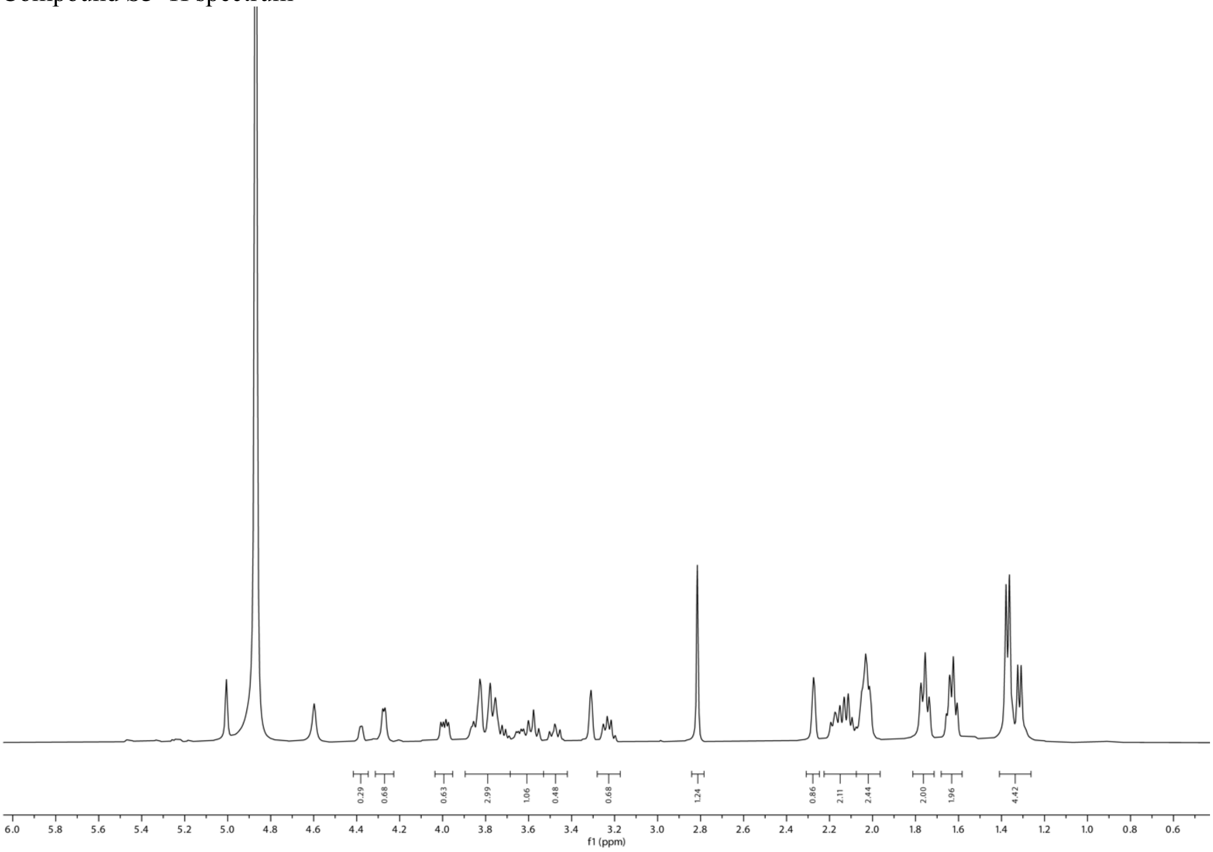


Compound **S5** $^{13}C$ spectrum

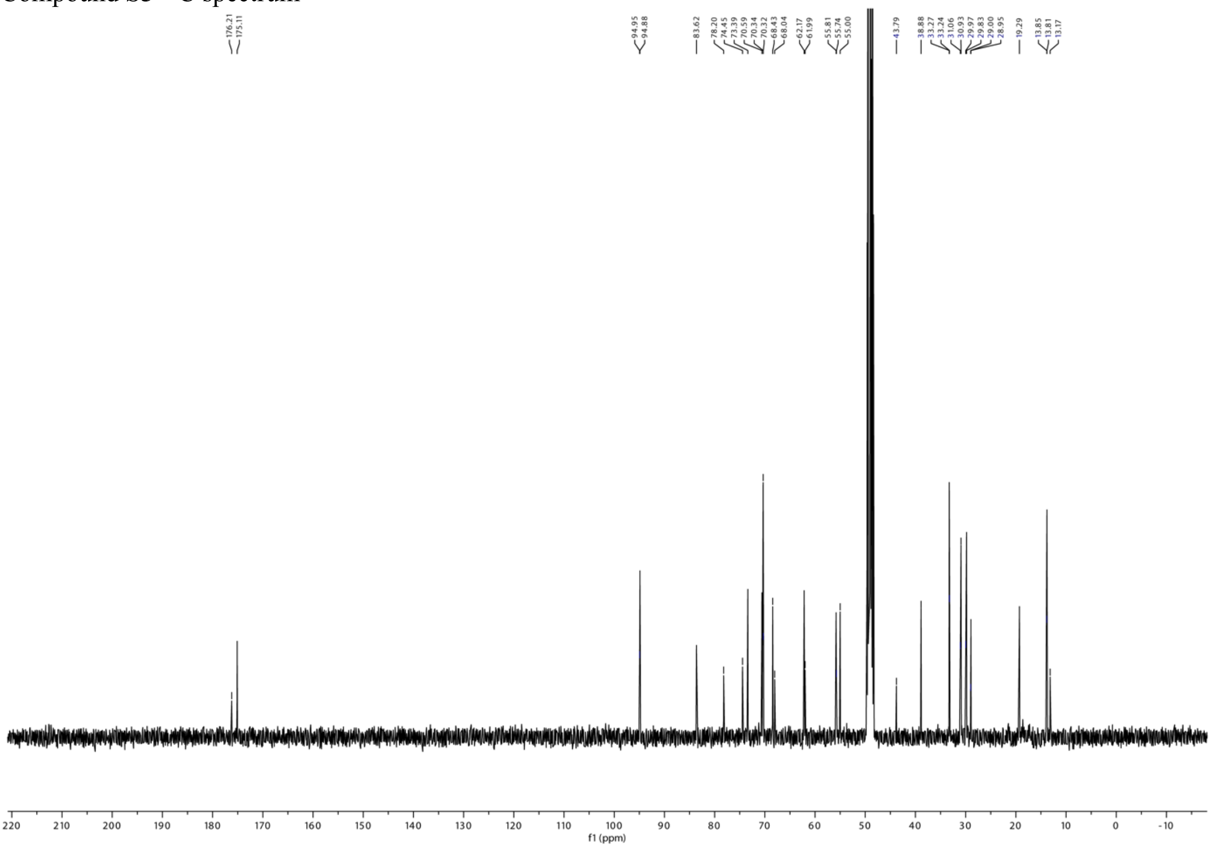

Compound **4** $^{1}H$ spectrum

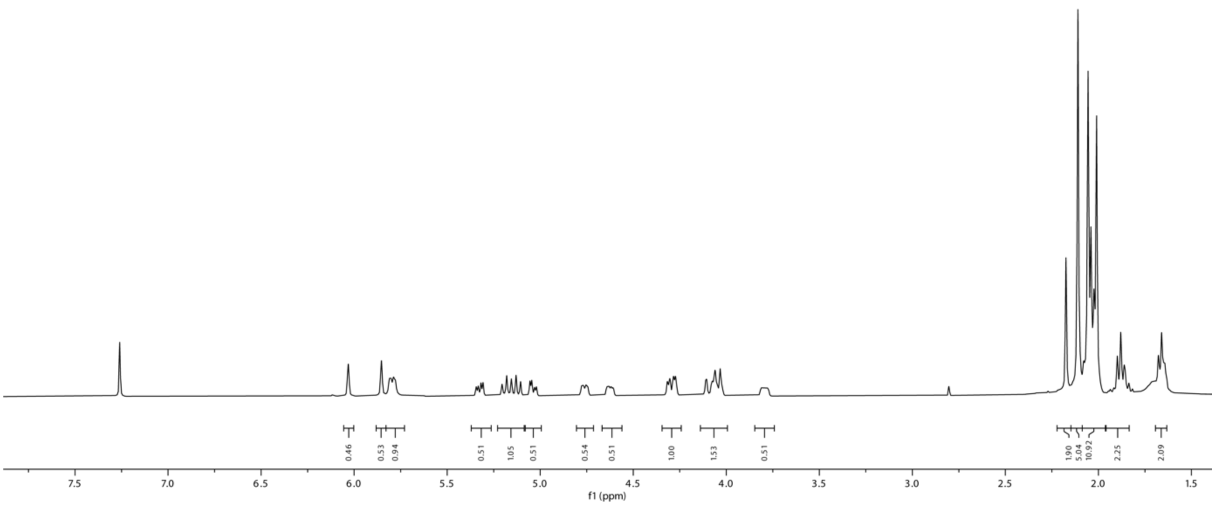

Compound **4** $^{13}C$ spectrum

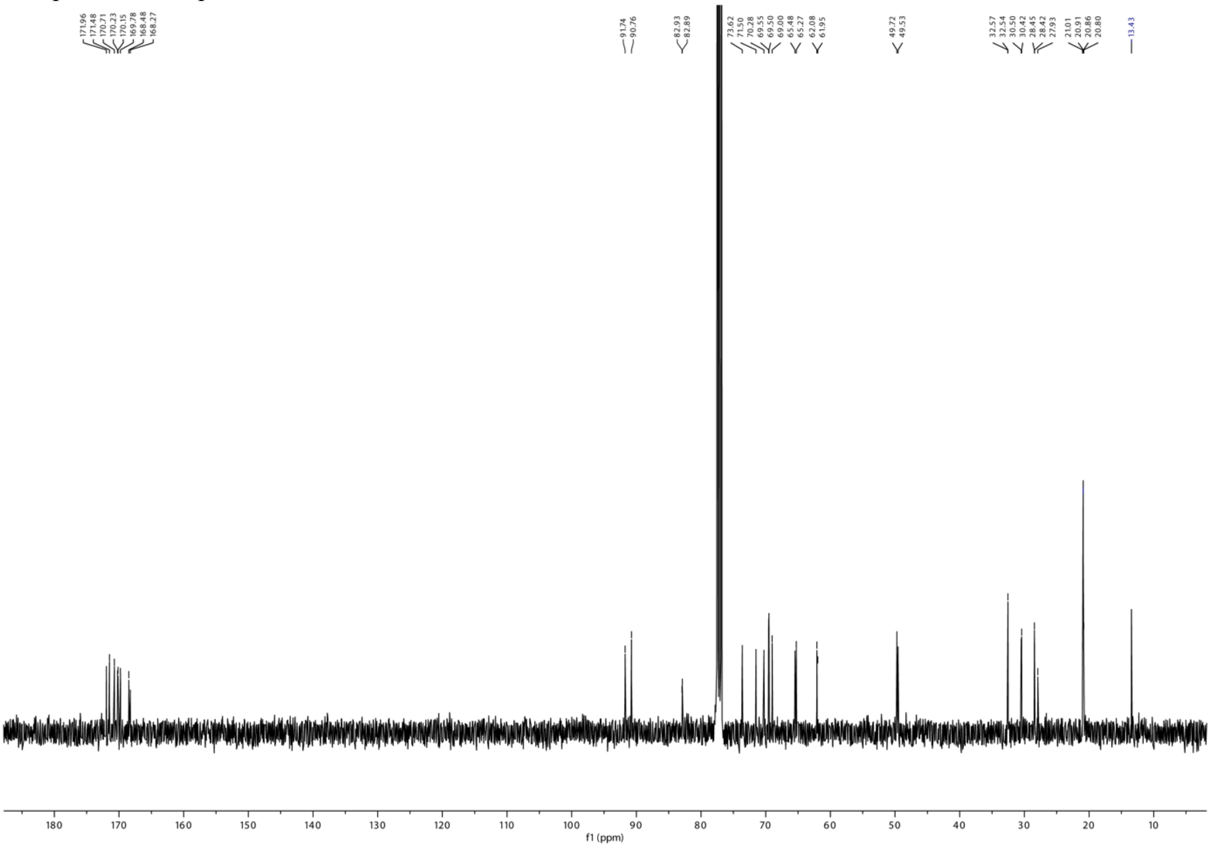
